# Intraspecific diversity in thermal performance determines phytoplankton ecological niche

**DOI:** 10.1101/2024.02.14.580366

**Authors:** Arianna I. Krinos, Sara K. Shapiro, Weixuan Li, Sheean T. Haley, Sonya T. Dyhrman, Stephanie Dutkiewicz, Michael J. Follows, Harriet Alexander

## Abstract

Temperature has a primary influence on phytoplankton physiology and affects biodiversity and ecology. To examine how intraspecific diversity and temperature shape plankton populations, we grew 12 strains of the ecologically-important coccolithophore *Gephyrocapsa huxleyi* isolated from regions of different temperature for ∼45 generations (2 months), each at 6-8 temperatures, and characterized the acclimated thermal response curve of each strain. Even with virtually identical temperature optima and overlapping cell size, strain growth rates varied between 0.45 and 1 day^-1^. While some thermal curves were effectively symmetrical, others had more slowly declining growth rates above the “thermal optimum,” and thermal niche widths varied between 16.7 and 24.8 °C. This suggests that different strains use distinct thermal response mechanisms. We investigated the ecological implications of such intraspecific diversity on thermal response using an ocean ecosystem simulation resolving distinct phytoplankton thermal phenotypes. Resolving model analogs of thermal “generalists” and “specialists” (similar to those observed in *G. huxleyi)* resulted in a distinctive global biogeography of preferred thermal niche widths with a nonlinear latitudinal pattern. We leveraged the model output to predict the ranges of the 12 strains we studied in the laboratory and demonstrated how this approach could refine predictions of phytoplankton thermal geographic range *in situ*. Our combination of observed thermal traits and modeled biogeography highlights the capacity of diverse groups to persist through temperature shifts.

**Significance Statement:** Intraspecific diversity in the phytoplankton may underpin their distribution. We show that within a single coccolithophore species, thermal response curves have diverse trait parameters. For example, many strains had a variable range of temperatures at which they could survive (thermal niche width). Adding this thermal niche width diversity to an ecosystem model simulation impacted phytoplankton coexistence and overall biomass. These observations show that thermal niche width is a gap in phytoplankton representation in ecosystem models that impacts modeled phytoplankton biogeography and concomitant carbon cycle dynamics. Including thermal tolerance is crucial to predictive modeling as ocean temperature dynamics change.

## Introduction

Temperature critically influences organism size (1–3), development (4), distribution (5, 6), and metabolic rate (7). Many organisms are reliant on environmental temperature, from microorganisms (8, 9), to corals (10, 11), and fish (12). The strong relationship between physiology and temperature indicates that higher average ocean water temperature will impact the abundance and distribution of species (13–16). Climate change is also expected to increase temperature variability (17, 18), which impacts organisms proportionally to their tolerable thermal range (19). Increasing climate variability and extremes are predicted to shape ecology, including phenology, species interactions, and dominant community assemblages (20–24).

Phytoplankton are vital to global primary production and carbon biogeochemistry (25). Their abundance and distribution correlates with temperature (26), and thermal response has multiplicative interactions with other drivers of phytoplankton fitness (27, 28). Thermal reaction norms describe the effect of temperature on growth rate, are measured for individual taxa in the laboratory, and indicate potential evolutionary tradeoffs between adaptive thermal mechanisms (such as the use of proteins with different temperature sensitivities (29), the rate or regulation of resource use (30, 31) (32)) and fitness (33).

The deviation of thermal reaction norms of individual ecotypes from a “standard” form is suggested to be among the most important markers of the role of phytoplankton diversity in shaping thermal response (34–36), but its ecological implications have not been explored in ecosystem models to date. The width of a thermal reaction norm indicates the tradeoff between being a temperature “generalist” capable of growing successfully over a wide temperature range, or a “specialist” with a narrower temperature range, but potentially with a growth rate advantage (33). The commonly used Eppley-Norberg model (37) parameterizes thermal reaction norm width explicitly (37–39). Thermal reaction norm width has broad relevance to ecology and species biogeography. Janzen’s rule (40, 41) hypothesizes that at higher latitude and elevation, higher thermal variability leads to wider thermal niche widths. Past work has found that phytoplankton frequently do not adhere to this expectation that thermal width increases with latitude (42). The diversity of phytoplankton makes it difficult to extrapolate point measurements of thermal width and local temperature to community trends, which is key to interpreting phytoplankton studies in the context of ecological theory.

To directly interrogate the effect of diversity on thermal response, we used G*ephyrocapsa* (formerly *Emiliania*) *huxleyi* as a model system. *G. huxleyi* is frequently grown in the laboratory as a model phytoplankter due to its global distribution and ease of culturing (43) as well as its developed molecular resources (44). Moreover, intraspecific diversity within *G*. *huxleyi* is well-described, including in calcite elemental composition, growth rate, C:N ratio, and thermal response (43, 45–48). Distinct *G. huxleyi* isolates have unique thermal reaction norms (36, 49), and strain diversity can shift thermal reaction norms in ecologically significant ways (50). Strain identity thus may determine viral resistance via lipid remodeling (51), alkenone composition (47), elemental composition and change (52), and trace metal speciation and use (53–55), all of which may determine both phytoplankton community composition and nutrient export. The *G. huxleyi* species complex is unusually resilient to temperature among coccolithophores (56), which may further increase its importance as global mean temperatures rise. *G. huxleyi* also has a level of diversity appropriate to test hypotheses about coccolithophore and general phytoplankton thermal physiology. Coccolithophores are frequently cited as suited to low temperature, low turbulence, oligotrophic conditions, which drives predictions of declines in this group as a consequence of climate warming (57). Despite this prediction, *G. huxleyi* expansions have recently been observed *in situ* (58, 59). There is a pressing need to examine undersampled thermal niche traits in *G. huxleyi* and other phytoplankton, as niche width and strain diversity could impact group coexistence and overall biomass and hence explain unexpected trends in the success of competing phytoplankton taxa.

Here, we selected 12 globally-distributed *G. huxleyi* isolates to assess the degree of thermal response curve variability across strains. To explore the impact of observed thermal trait diversity on total biomass and thermal type coexistence, we designed a model simulation for a general phytoplankter with varying thermal optima (following (60)) and included variable thermal response norm width among model “ecotypes”. The large, relatively geographically-balanced collection of *G. huxleyi* strains that we collected expanded available thermal response data and provided sufficient resolution to implement a diversity-resolving model simulation. We combined the model simulation with the laboratory dataset to demonstrate that model output can predict and diagnose reasons for the success of dominant traits across ocean regions. Our study bridges increasing recognition of the importance of strain-specific processes with the impacts of intraspecific diversity on typical model representations of thermal response and global thermal range predictions.

## Results and Discussion

### *G. huxleyi* thermal reaction norms are intraspecifically variable

To quantify the intraspecific variability of phytoplankton thermal response due to local thermal habitat, we selected 12 strains of *G. huxleyi* isolated from across the global ocean and from a variety of global environmental regimes (Figure 1). We acclimated strains to temperatures in the lab for at least 2 months and then characterized thermal reaction norms. The 12 strains of *G. huxleyi* in this study have distinct maximum growth rate, ratio between maximum growth rate and optimum temperature, and thermal range (Figure 2). The total range in measured thermal widths was 8.1°C, the range in thermal optimum was 11.4°C, and the range between maximum growth rates at the thermal optima of the strains was 0.69 day^-1^. We found no significant relationship between thermal optimum and the growth rate at the thermal optimum (Kendall’s tau: T=34, tau=0.03, *p*=0.95). These results indicate that the strains we measured did not follow any straightforward scaling between average preferred temperature and either thermal range or maximum growth rate, highlighting the ecosystem relevance of intraspecific diversity under constant ambient environmental conditions. The 12 *G. huxleyi* strains examined here also had a broader range in thermal optimum than previously observed (48, 49, 57) due largely to our addition of a strain from the Southern Ocean (RCC6071) . The data from the 12 strains we examined reaffirmed that coccolithophores have a wide range in maximum growth rates within a relatively small range of thermal optima (Figure 2B). The Southern Ocean strain also has a higher maximum growth rate than several strains with higher optimum growth temperatures (Figure 2B).

**Figure 1.**
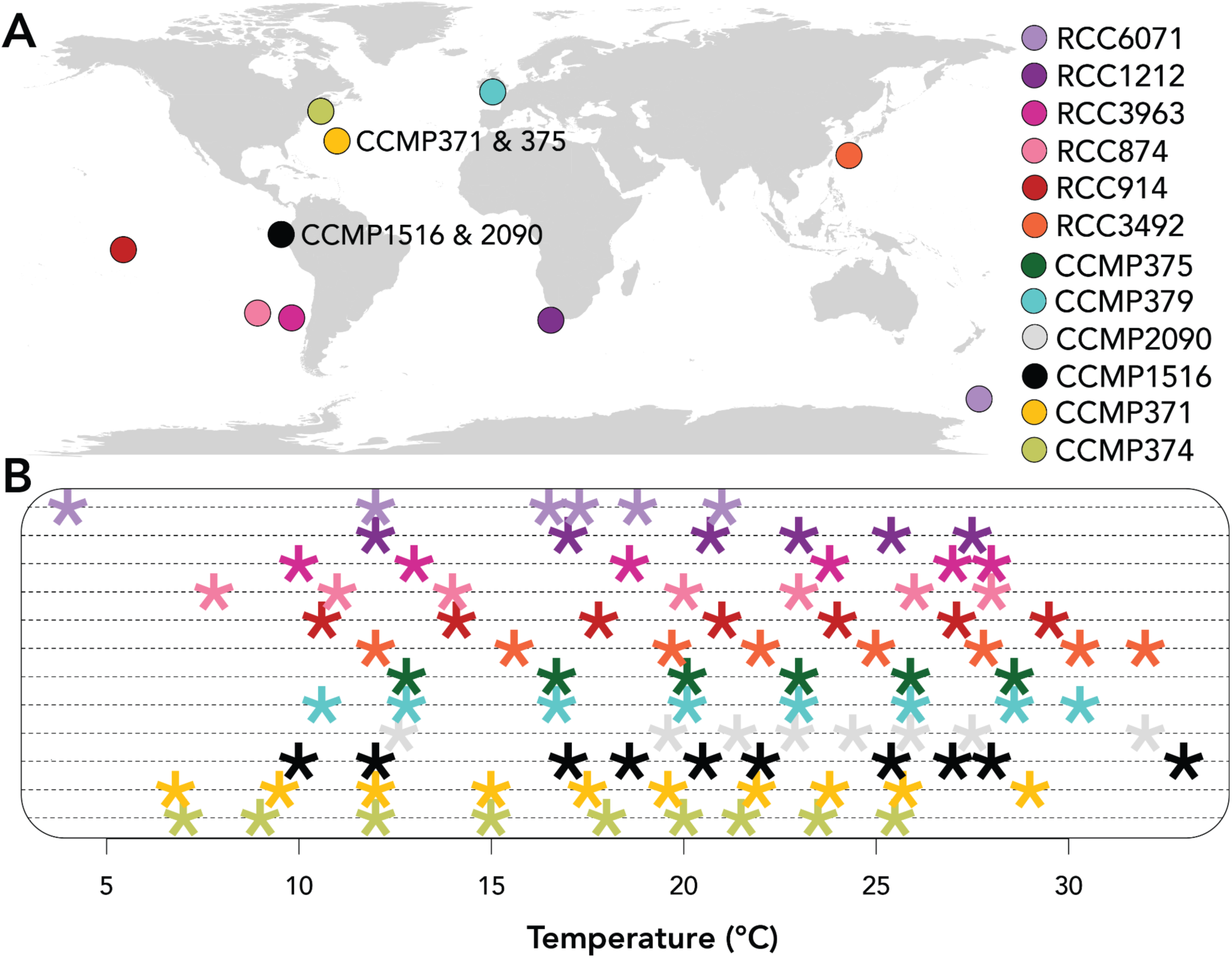
Isolation location and measurement frequency of strains of *Gephyrocapsa huxleyi* tested. A: Location of original isolation for each strain, which were retrieved from either the Roscoff Culture Collection (RCC) or the National Center for Marine Algae and Microbiota (NCMA). B: Temperatures evaluated for each of the 12 strains.

**Figure 2.**
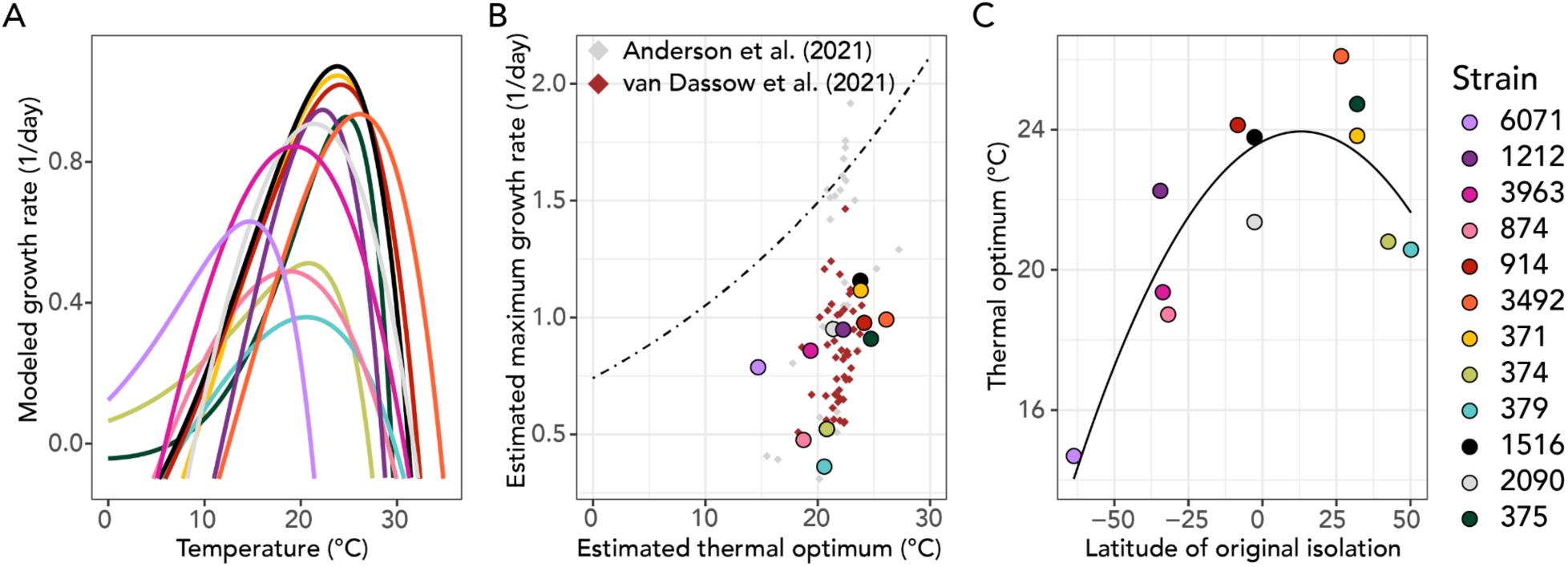
Thermal response parameters for the 12 tested strains of *Gephyrocapsa huxleyi*. A: Thermal response curves as parameterized by the Norberg equation for each strain. B: Maximum growth rate compared to the estimated thermal optimum for each strain; the size of each point indicates its thermal width. The black dotted line indicates the hypothesized Eppley relationship for coccolithophores from Anderson *et al*. (2021). C: Estimated thermal optimum by latitude of isolation, where strains with higher latitude of original isolation had higher thermal optimum values.

With the specific goal of identifying generalists and specialists, we compared our thermal performance characterizations to a prior data compilation for coccolithophores (57), and added a recent set of thermal characterizations of *G. huxleyi* (*49*) (Figure 2B). Our globally-sampled data affirm that thermal trait diversity within the cosmopolitan species *G. huxleyi* is representative of all known thermal response data for coccolithophores (Figure 2A,B; Supplementary Figure 1), hence *G. huxleyi* is exceptionally phenotypically variable. However, the total diversity of coccolithophore thermal responses is undersampled–only four other species of coccolithophore were present in the two datasets combined (49, 57). Our work underscores the importance of considering intraspecific variability, in that characterizing a single isolate is insufficient to capture the flexibility of phytoplankton thermal response and inform parameterizations of phytoplankton thermal limits in models (61–63).

Generalist and specialist strategies may influence the ecology of phytoplankton within and between thermal types and determine the water temperatures at which they can be successful. We evaluated *G. huxleyi*’s thermal niche width in the laboratory experiments by recalculating the width of the thermal response curve using the temperatures at which the simulated growth rate via the Norberg parameterization crossed zero. The 12 strains fell along a range of thermal width values, with strain RCC1212 being the most “specialist”, while strain RCC3963 was the most “generalist” (Figure 2B). Given the relationship between maximum growth rate and thermal optimum and other factors influencing growth, it is difficult to ascertain from laboratory data whether a penalty exists. We could not assign a straightforward growth rate cost to a broader thermal niche width. Although more thermal range flexibility would likely necessitate a lower maximum growth rate (33, 64), neither our data nor previous data compilations supported a significant penalty (Supplementary Figure 25). The relationship between maximum growth rate and thermal optimum complicates the evaluation of the cost.

We used a second thermal width parameter for the range of temperatures that fell within 80% of the maximum measured growth rate. We call this the “plateau parameter,” since it captures the scenario that we frequently observed in which a range of temperatures appeared to be close to equally suitable for growth (Supplementary Figure 2). The plateau parameter had a range of 6.5°C between strains, and similarly had no significant relationship with optimum temperature (Kendall’s tau: T=22, tau=-0.33, *p*=0.15) or maximum growth rate at the thermal optimum (Kendall’s tau: T=25, tau=-0.24, *p*=0.31). The plateau parameter was weakly correlated with the thermal width (Kendall’s tau: T=47, tau=0.42, *p*=0.063). Strains with a high range of survivable temperatures tended to have high growth rates across that survivable temperature range, but their maximum growth rates did not occupy a uniform proportion of the range (Supplementary Figure 2). A larger measured thermal niche width only partially explained the larger range of temperatures around the thermal optimum with similar, near-maximum growth rates. The phytoplankton thermal types we measured were hence diverse in thermal optimum, thermal niche width, and range of temperatures with high, near-maximum growth rates.

### Varying modeled thermal width traits predicts distinct biogeographies of generalists and specialists

Our observations of the thermal traits of *G. huxleyi* highlight the variability in thermal niche width present in a single marine species. To understand how this diversity may impact *G. huxleyi* and other phytoplankton distribution at the ecosystem scale, we took an ecological modeling approach. To isolate the role of thermal reaction norm in determining phytoplankton distribution and biomass, we used the Darwin model, an ecosystem layer on the MIT general circulation model (60, 65). We resolved sixty identical phytoplankton ecotypes with the same light and nutrient requirements and susceptibility to predation. These ecotypes only differed in their thermal optima and thermal niche widths. We chose six thermal widths that linearly spanned the approximate range of thermal niche widths we observed across *G. huxleyi* strains (Figure 3C). Both our experiments and a previous data compilation did not find a clear relationship between thermal niche width and growth rate (42), but imposing a hypothesized growth rate penalty in the model ensured that a wider niche width would not be a universally beneficial trait (Supplementary Figure 4). To interpret the impact of this penalty, we compared our results to a no and a high penalty model scenario (Supplementary Information; Supplementary Figure 4. Comparing the no- and high-penalty scenario to the intermediate penalty we used in this study revealed that while generalist biomass correlates strongly with temperature variability in the no-cost scenario (Supplementary Figure 4A), there was low overall thermal niche width diversity, which contrasts with our laboratory observations. The high-cost scenario was completely specialist-dominated and showed the least correlation to temperature variability (Supplementary Figure 4C). The intermediate penalty scheme that we adopted provided a reasonable tradeoff between these two extremes that was consistent with the observation that many different thermal niche widths are indeed found *in situ*, suggesting that a penalty may exist even if not directly observed.

**Figure 3.**
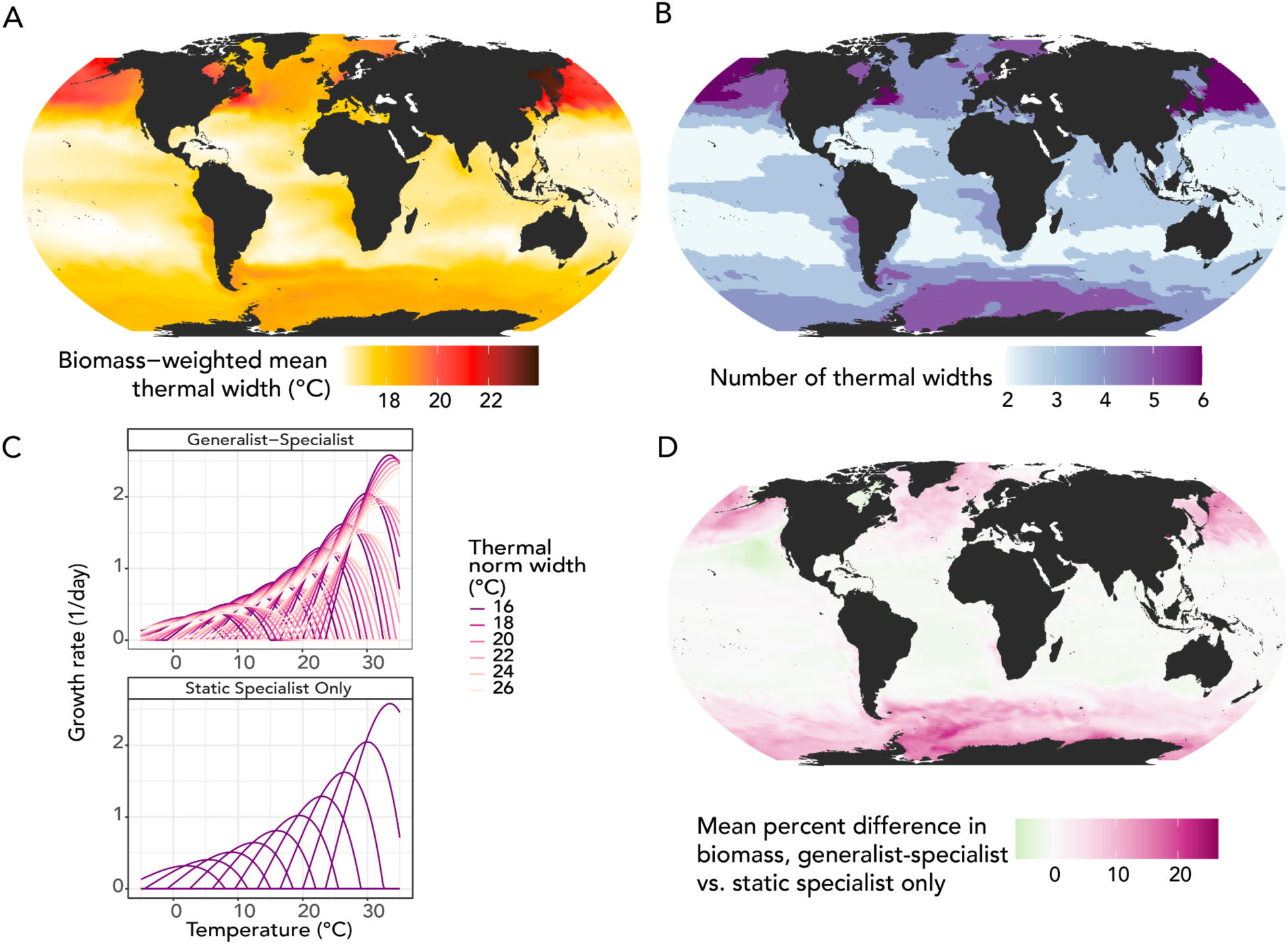
Thermal niche width modifications to the Darwin model simulation reveal distinct global biogeography of phytoplankton generalists and specialists. A: Mean biomass-weighted thermal niche width of 60 phytoplankton functional types in the Darwin model simulation. Whereas theory suggests that specialists should dominate in high-latitude regions with high seasonality, we found that specialists tended to play a larger role in the total phytoplankton community in the subtropical gyres, whereas generalists with higher thermal niche widths thrived in temperate, mid-latitude regions. B: Number of thermal niche widths observed with at least 1% biomass in the generalist-specialist simulation; more purple colors indicate more thermal niche widths present, up to a maximum of 6. C: Static thermal niche width (left) vs. specialist-generalist (right) thermal reaction norms; each color corresponds to a different thermal niche width, with lighter colors indicating a more generalist thermal niche width. D: Percentage difference in mean modeled biomass in the final year of the simulation between the generalist-specialist simulation and the static specialist only simulation. Darker pink colors indicate higher biomass in the generalist-specialist simulation, whereas green colors indicate higher biomass in the static specialist only simulation.

The largest relative populations of generalists were found in the north Pacific Ocean and northwest Atlantic (Figure 3A), which were areas of high temperature variation in the model (Supplementary Figure 3; Figure 3). However, some regions that favored generalists, such as areas of the Southern Ocean, did not have high thermal variability. Further, regions enriched with specialists included oligotrophic gyres with low thermal variability (Figure 3A). The observation that regions with higher thermal variability tend to favor generalists in our model is partially compatible with generalizations of Janzen’s hypothesis (40, 41). However, our model reveals that physical features or the timing of seasonal changes in temperature and nutrient availability may result in an outsized role of resource availability (66) over thermal variability in some regions (*e.g.*, Supplementary Figures 4 and 5).

In this intermediate cost simulation, mean thermal niche width and number of coexisting thermal niche widths (Figure 3A,B) corresponded to ocean biogeochemical provinces, such as oligotrophic gyres and boundary currents. Due in part to these features, abundance patterns of individual thermal types did not necessarily correlate to thermal variability via standard metrics. These features may explain unexpected observations in thermal niche width relative to latitude or local temperature. For example, past work found that strains of *G. huxleyi* isolated from Bergen, Norway, a region with comparatively high thermal variability, actually had a higher mean thermal reaction norm width than isolates from Azores, Portugal, where thermal variability was lower (36). The model predicts approximately the same mean thermal niche width (Figure 3A; Supplementary Figure 5), and coexistence between generalists and specialists (Supplementary Figure 8) at these two locations. This result indicates the flexibility of the model to predicting thermal niche width coexistence amid differences in temperature variability. While regions of very high thermal variability in the model had correspondingly high average thermal niche widths, average weighted thermal niche width more frequently corresponded to basin biogeography–hence local resource availability (66–69)–than absolute range or standard deviation in environmental temperature (Supplementary Figures 4, 5, 6). This could be due to geographic separation between basins, advection of specific water masses, nutrient availability, or physical mixing.

To further explore the benefit of being a generalist, we compared the simulation with both specialists and generalists (generalist-specialist experiment) to one with only specialists (Figure 3C). We found that total biomass was generally lower in the specialist-only simulation (Figure 3D). The largest changes in biomass occurred in generalist-dominated regions (Figure 3A,D). However, there were also some increases in biomass in regions that favored specialists, suggesting that generalists were in fact less productive in these regions, but had nevertheless persisted in the generalist-specialist experiment. Hence, biogeochemical conditions promoted the existence of regional niches selecting for diverse thermal widths over maximized community growth rate. Notably, while many regions that had more generalists than specialists had higher total biomass in the generalist-specialist simulation as compared to the specialist-only simulation, change in biomass was not proportional to the relative abundance of generalists (Figures 3A,D). This suggests a higher relative importance of other factors (*e.g*., nutrients or phenology) in these regions.

We also compared the specialist-only simulation to a generalist-only simulation, which revealed different global patterns in the benefit of being a specialist versus a generalist (Supplementary Figures 9 and 10). Most of the global ocean had higher biomass in the generalist-only simulation. However, regions like the northwest Pacific and Sargasso Sea had unexpectedly higher biomass in the specialist-only scenario (Supplementary Figures 9 and 10) despite high thermal variability (Supplementary Figure 3). Short-term bursts in specialist biomass are only beneficial when fully coupled to the timescale of temperature change, hence mismatch between these two timescales may explain difference in the specialist-benefit tradeoff between ocean regions. Expected impacts differed between the generalist-only and the generalist-specialist simulation, indicating that the presence of both specialists and generalists influences overall biomass. The simulation results reinforced the observation that thermal niche width influences overall predicted biomass in an ecosystem model simulation, despite not always correlating directly with local temperature.

### Strain geographic predictions highlight the role of intraspecific diversity in biogeography

The ecosystem model simulation output can be leveraged to determine which thermal traits are likely to be most successful in each ocean region. We used the ecosystem model output to predict the distribution of the twelve strains of *G. huxleyi* that we measured in the laboratory (Figure 4). Using the minimum and maximum temperatures of the model thermal types that could survive at each latitude and longitude, we calculated the probability that a hypothetical “thermal type” with the same thermal maximum and minimum of each of the measured laboratory strains could persist. We found agreement between the location of original isolation of the strain and the predicted model probability that the strain would exist in that location. For example, 6 of the strains had greater than 15% probability and 3 had greater than 50% probability (Supplementary Figure 11). The 3 strains with less than 1% probability of existence in the model grid point corresponding to their isolation location (RCC6071, RCC914, and CCMP1516) had high probability values nearby (Supplementary Figure 11). Latent diversity within *G. huxleyi* indicates the ability of the species to survive across virtually all global ocean regimes, yet no individual strain tested had a greater than 1% probability in all simulated ocean regions (Supplementary Figure 12). This result indicates that intraspecific variability in thermal trait diversity can increase the overall success of a species in surviving a range of temperatures and may be a mechanism that underpins the co-existence of strains *in situ*.

**Figure 4.**
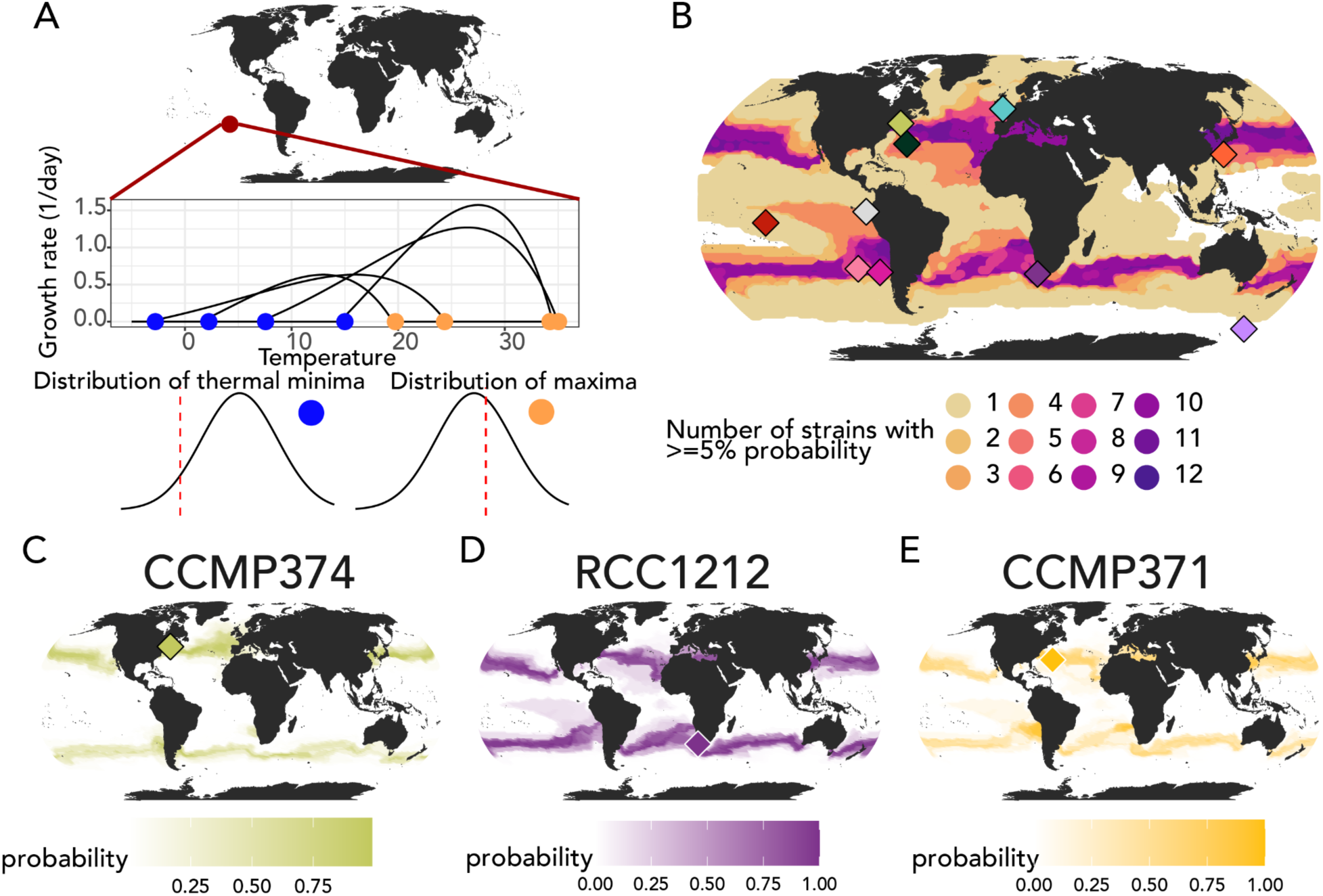
Probabilistic projections of the environmental distributions of each of the strains of *Gephyrocapsa huxleyi* measured in the laboratory. **A**: A representation of the methodology used to calculate probabilities; a distribution was built from the thermal minima and maxima of all strains that existed at each gridded latitude and longitude point in the model and constituted at least 1% of the population. Probability was calculated using the probability density function for each distribution, then the two probabilities were multiplied. **B**: The number of strains in each latitude and longitude point with at least a 5% probability of occurring from 0 (white) to 12 (dark purple); colored diamonds indicate the isolation locations of each of the 12 strains and are identical to the gridded locations and colors of Figure 1A. **C**: Predicted probability using the method from 4A for strain CCMP374. **D**: Predicted probability using the method from 4A for strain RCC1212. E: Predicted probability using the method from 4A for strain CCMP371.

### Implications for modeled current and future phytoplankton habitat

Taken together, the model analysis and laboratory results demonstrate that the width of the thermal response curve can have a strongly deterministic influence on the simulated distribution of phytoplankton in a diversity-resolving ecosystem model. The varied thermal niche widths of the strains we measured in the laboratory suggest that strains use different mechanisms to cope with environmental temperature. Genome sequencing paired with gene expression studies will illuminate what specific differences in biological pathways may be responsible for different observed thermal parameters between strains (70). Thermal response mechanisms may also have variable resource reliance, which could reduce, eliminate, or augment each strain’s thermal response (71–73). Future work on the thermal sensitivity of phytoplankton to other environmental drivers (*e.g*., nutrient-temperature relationships and trace metal availability) should also consider intraspecific variation. Our observation that generalist-specialist simulations show greater biomass in thermally-variable ocean regions may impact predicted phytoplankton resilience to future climate, including increasing thermal variability (17, 20, 74) and marine heat waves (75). For example, coccolithophores may be able to persist during short-term heat waves via their slowly increasing maximum growth rates at their optimum temperatures and diversity of thermal niche widths, which may increase stored resources and resilience to temperature fluctuation (76, 77). Because *G. huxleyi* tends to be comparatively resilient to warming temperatures and fluctuating nutrients among coccolithophores (56, 78), the intraspecific diversity in thermal niche (*e.g*., generalist v. specialist) observed here may offer a still greater mechanistic advantage against temperature warming and variability. Measuring the phenotypic flexibility that confers diverse thermal traits and encoding it in models is hence essential to projecting future change in the balance of phytoplankton functional types with global climate.

## Materials and Methods

### Laboratory culturing

Starter cultures were obtained from the Bigelow Laboratory for Ocean Science’s National Center for Marine Algae and Microbiota (NCMA) for strains CCMP371, 374, 375, 379, and 2090, and from the Roscoff Culture Collection (RCC) for strains RCC874, 914, 1212, 3492, 3963, 4567, and 6071. Strain CCMP1516, which is a descendant of the same original isolation as Strain 2090, was obtained from the Dyhrman laboratory at Columbia University and has been observed to calcify, whereas strain CCMP2090 no longer calcifies in culture. Maintenance cultures of each of the thirteen strains were kept at 18°C under a 14:10 light/dark cycle and transferred approximately once per month. Four fluorescent tube bulbs were positioned below the culture vials, one per row of the thermal block, such that each set of culture tubes was situated directly above a bulb. Approximately 10 centimeters from the light source, a light level of approximately 24 umol m^-2^ s^-1^ was measured. Natural seawater from Vineyard Sound, MA, USA was used as the base for all media recipes. Strains were maintained in standard L1 media without silica (reported herein as L1-Si; as per https://ncma.bigelow.org/algae-media-recipes). Strains CCMP1516 and 371 were maintained in low nutrient media to help retain their calcification state, where all nutrients, trace metals, and vitamins were added at 1/25th of the standard concentration (reported herein as L1/25-Si; as per https://ncma.bigelow.org/algae-media-recipes).

An aluminum thermal block was used to achieve a thermal gradient over which strains could be incubated at increasing temperatures. The aluminum block has 80 openings across 4 rows and 20 columns for 25 mm diameter glass culture tubes. Thermal equilibrium is maintained using insulation, and a shared light source at the bottom of the thermal block enables constant light levels across the experiment (79, 80). A circulating water bath was used to keep one side of the thermal gradient cool, while a heating element on the other side set the maximum temperature of the experiment; the low and high temperatures were selected for each set of measured strains based on their expected thermal tolerance. Temperature extremes were set according to realistic temperatures for the strain being tested, and temperatures were measured and recorded regularly throughout the experiments.

Each strain was transferred via 1 mL aliquots from the maintenance stock to 6-7 different temperatures spanning the thermal gradient at the beginning of the experiment. Strains CCMP1516 and 371, as well as strain CCMP2090 for inter-comparability with descendant of the same lineage strain CCMP1516, were transferred into and maintained in the L1/25-Si media, while the remainder of the strains were transferred into and maintained in L1-Si media for all subsequent transfers in the thermal experiment. Cell abundances and optical properties were measured daily using flow cytometry, with either a Guava easyCyte HT 2 or 3 (Luminex, USA) (Strains RCC874, CCMP371, CCMP379, and CCMP374) or a BD Accuri C6 Plus equipped with a C-Sampler (BD, USA) (Strains RCC1212, RCC6071, CCMP1516, CCMP2090, RCC3963, RCC914, RCC3492, and CCMP375). Each strain population was maintained in triplicate rows with identical temperatures in semi-continuous culture for a minimum of 45 generations, or approximately 2 months. Data were manually curated to ensure that compatible points in the culture cycle were used for growth rate calculations. Maximum growth rates were computed using the last recorded time point in exponential phase and the first recorded time point in exponential phase according to the equation:

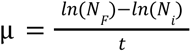

where *N_f_* is the final recorded concentration in exponential phase as measured in cells per milliliter, *N_i_* is the first recorded concentration, and *t* is the duration in days. The final growth rate is expressed in dimensionless units of per day. Final thermal reaction norms were constructed using growth rates computed from up to 3 semi-continuous transfers when data meeting the minimum quality thresholds were available.

### Thermal response curve parameterization

We used the equation from Norberg (2004) to parameterize phytoplankton growth rates across the strains we studied:

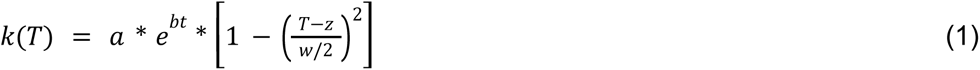

where *T* is temperature, *k(T)* is specific growth rate, *w* is the estimated thermal niche width, and *a*, *b*, and *z* are opaque shape parameters, commonly fit using methods like maximum likelihood estimation (39, 81, 82). We used the bbmle package (version 1.0.25) (83) to estimate parameter values for this equation for each of the strains. We estimated the thermal optimum from the equation by using the optimize function in R with the maximum parameter set over an interval in temperatures from zero to 40°C. We recalculated the thermal width for the data from (57) using 0.01 as a threshold value for when growth rate was zero at the intersection points, since the parameterizations tended to have unrealistically high reported thermal widths when the estimated thermal performance curves had long tails with near-zero growth rates.

We calculated the “plateau parameter” (the range of temperatures over which measured growth rate was within 80% of the maximum measured growth rate) using the same search procedure to determine the intersection points. The code used to calculate these quantities is available in the published GitHub repository.

### Parameterization of Darwin 3-dimensional model simulation

To simulate more diverse thermal response curve shapes, we modified the Darwin model (60, 65, 84). The Darwin model is designed to be flexible in the number and types of plankton to include. Here to focus on the relevance of thermal norm structure alone, we implement a setup where the phytoplankton types we include (either 10 or 60 types) are identical to each other except for their thermal norm and are grazed equally by the single zooplankton grazer. The 10 or 60 phytoplankton functional types had one of 10 thermal optimum values. We compared two simulation configurations: thermal optimum only, wherein only thermal optimum varied between 10 different types and roughly linearly spaced optimum values varied between 0°C and 31.5°C corresponding to typical thermal optimum values for coccolithophores. In a second model configuration, the same 10 thermal optimum types were simulated between 0°C and 31.5°C, but each thermal optimum type also corresponded to 6 different thermal niche width values (16, 18, 20, 22, 24, and 26°C). These thermal niche width values each corresponded to a different value of both the parameters *a* and *w* in the Norberg curve parameterization of the thermal curve. We penalized the *a* parameter of each functional type by 0.05 for every degree Celsius wider its width was than the baseline specialist phytoplankton functional type. This imposed a cost to adopting the generalist lifestyle in the model. Because our strains varied in thermal niche width in the laboratory without predictable changes in maximum growth rate, we applied a uniform penalty rather than using an observed laboratory relationship between thermal width and maximum growth rate. A single grazer class was simulated in the model, with grazing rate scaled by temperature according to the expression:

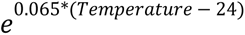

In each of these two experiments, the phytoplankton types were initialized with identical biomass and the simulation was run for 10 years. The biogeography of the different phytoplankton types reaches a quasi-steady seasonal cycle after about 3 years of integration. We present results from the last year of this simulation.

### Analysis of Darwin model output

The Darwin model results were saved in time-averaged intervals of one month, and then the surface layer of the simulation was extracted from the model output. Biomass was measured in the model in units of mg/m^3^ in carbon units for each phytoplankton type. Biomass-weighted thermal niche width was calculated at each gridded latitude and longitude location according to the expression:

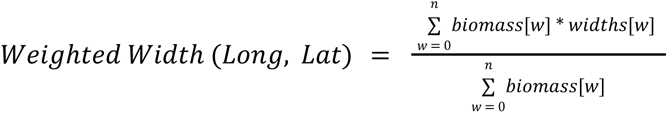

All maps were generated using the ggalt package (85) and ggplot within the R statistical computing environment (version 4.1) (86, 87). The map projection was produced using the coord_proj function and parameters “+proj=robin +lon_0=0 +x_0=0 +y_0=0 +ellps=WGS84 +datum=WGS84 +units=m +no_defs”. RColorBrewer was used to create gradient color fills for maps (88).

Presence or absence of thermal niche widths was estimated by considering widths to be “present” when they constituted at least 1% of total community biomass in at least one time-averaged month of the simulation.

### Projection of thermal habitat of laboratory strains

The maximum thermal habitat of each strain was calculated by taking all strains that had ever constituted at least 1% of the community at each latitude and longitude point. The maximum and minimum temperature of each strain as assessed by the thermal width was extracted and used to build two different normal distributions for each latitude and longitude point. For each of the twelve strains, the probability density function value was calculated for the minimum and maximum temperature values observed for the strain, then normalized to the probability of the mean of the distribution (such that an observed thermal minimum/maximum matching the mean of the distribution would have a probability of 1). The two probabilities were combined by multiplication. We assumed that the two probabilities were independent and could be combined in this way because we allowed the thermal niche width and optimum to vary. Hence, the observed minimum temperature value could not be used to predict the mean of the distribution of maximum temperature values observed in the model and the two distributions can be assumed independent.

## Data Sharing Statement

All code used to create figures is available on GitHub at https://github.com/AlexanderLabWHOI/2024-Krinos-Ghux-Darwin. All growth rate data used in the model and the thermal parameters calculated for each strain is uploaded to the online GitHub repository as well as available in Supplementary Tables 1-3.

## References

1. J. Forster, A. G. Hirst, G. F. Esteban, Achieving temperature-size changes in a unicellular organism. ISME J. 7, 28–36 (2013).

2. M. Winder, U. Sommer, Phytoplankton response to a changing climate. Hydrobiologia 698, 5–16 (2012).

3. T. Zohary, G. Flaim, U. Sommer, Temperature and the size of freshwater phytoplankton. Hydrobiologia 848, 143–155 (2021).

4. J. F. Gillooly, E. L. Charnov, G. B. West, V. M. Savage, J. H. Brown, Effects of size and temperature on developmental time. Nature 417, 70–73 (2002).

5. G. N. Somero, Linking biogeography to physiology: Evolutionary and acclimatory adjustments of thermal limits. Front. Zool. 2, 1 (2005).

6. E. P. Jeffree, C. E. Jeffree, Temperature and the Biogeographical Distributions of Species. Funct. Ecol. 8, 640–650 (1994).

7. V. M. Savage, J. F. Gilloly, J. H. Brown, E. L. Charnov, Effects of body size and temperature on population growth. Am. Nat. 163, 429–441 (2004).

8. J. K. Jansson, K. S. Hofmockel, Soil microbiomes and climate change. Nat. Rev. Microbiol. 18, 35–46 (2020).

9. B. K. Singh, R. D. Bardgett, P. Smith, D. S. Reay, Microorganisms and climate change: terrestrial feedbacks and mitigation options. Nat. Rev. Microbiol. 8, 779–790 (2010).

10. R. Carballo-Bolaños, D. Soto, C. A. Chen, Thermal Stress and Resilience of Corals in a Climate-Changing World. J. Mar. Sci. Eng. 8, 15 (2019).

11. K. R. Bairos-Novak, M. O. Hoogenboom, M. J. H. van Oppen, S. R. Connolly, Coral adaptation to climate change: Meta-analysis reveals high heritability across multiple traits. Glob. Chang. Biol. 27, 5694–5710 (2021).

12. F. T. Dahlke, S. Wohlrab, M. Butzin, H.-O. Pörtner, Thermal bottlenecks in the life cycle define climate vulnerability of fish. Science 369, 65–70 (2020).

13. R. Cavicchioli, et al., Scientists’ warning to humanity: microorganisms and climate change. Nat. Rev. Microbiol. 17, 569–586 (2019).

14. M. P. Lesser, Coral reef bleaching and global climate change: can corals survive the next century? Proc. Natl. Acad. Sci. U. S. A. 104, 5259–5260 (2007).

15. R. Vaquer-Sunyer, C. M. Duarte, Temperature effects on oxygen thresholds for hypoxia in marine benthic organisms. Glob. Chang. Biol. 17, 1788–1797 (2011).

16. B. Abirami, M. Radhakrishnan, S. Kumaran, A. Wilson, Impacts of global warming on marine microbial communities. Sci. Total Environ. 791, 147905 (2021).

17. M. Rummukainen, Changes in climate and weather extremes in the 21st century. Wiley Interdiscip. Rev. Clim. Change 3, 115–129 (2012).

18. N. W. Arnell, J. A. Lowe, A. J. Challinor, T. J. Osborn, Global and regional impacts of climate change at different levels of global temperature increase. Clim. Change 155, 377–391 (2019).

19. P. K. Thornton, P. J. Ericksen, M. Herrero, A. J. Challinor, Climate variability and vulnerability to climate change: a review. Glob. Chang. Biol. 20, 3313–3328 (2014).

20. D. A. Vasseur, et al., Increased temperature variation poses a greater risk to species than climate warming. Proc. Biol. Sci. 281, 20132612 (2014).

21. C. Parmesan, G. Yohe, A globally coherent fingerprint of climate change impacts across natural systems. Nature 421, 37–42 (2003).

22. J. M. Tylianakis, R. K. Didham, J. Bascompte, D. A. Wardle, Global change and species interactions in terrestrial ecosystems. Ecol. Lett. 11, 1351–1363 (2008).

23. S. Lavergne, N. Mouquet, W. Thuiller, O. Ronce, Biodiversity and Climate Change: Integrating Evolutionary and Ecological Responses of Species and Communities. Annu. Rev. Ecol. Evol. Syst. 41, 321–350 (2010).

24. T. Mock, et al., Bridging the gap between omics and earth system science to better understand how environmental change impacts marine microbes. Glob. Chang. Biol. 22, 61–75 (2016).

25. J. D. H. Strickland, PHYTOPLANKTON AND MARINE PRIMARY PRODUCTION. Annual Reviews Microbiology (1965).

26. D. Righetti, M. Vogt, N. Gruber, A. Psomas, N. E. Zimmermann, Global pattern of phytoplankton diversity driven by temperature and environmental variability. Sci Adv 5, eaau6253 (2019).

27. R. W. Eppley, Temperature and phytoplankton growth in the sea. Fish. Bull. 70, 1063–1085 (1972).

28. G.-Y. Rhee, I. J. Gotham, The effect of environmental factors on phytoplankton growth: Temperature and the interactions of temperature with nutrient limitation1. Limnol. Oceanogr. 26, 635–648 (1981).

29. G. N. Somero, Proteins and temperature. Annu. Rev. Physiol. 57, 43–68 (1995).

30. P. A. Thompson, M.-X. Guo, P. J. Harrison, EFFECTS OF VARIATION IN TEMPERATURE. I. on THE BIOCHEMICAL COMPOSITION OF EIGHT SPECIES OF MARINE PHYTOPLANKTON^1^. J. Phycol. 28, 481–488 (1992).

31. F. A. Q. Sayegh, D. J. S. Montagnes, Temperature shifts induce intraspecific variation in microalgal production and biochemical composition. Bioresour. Technol. 102, 3007–3013 (2011).

32. K. G. Baker, R. J. Geider, Phytoplankton mortality in a changing thermal seascape. Glob. Chang. Biol. 27, 5253–5261 (2021).

33. M. J. Angilletta, R. S. Wilson, C. A. Navas, R. S. James, Tradeoffs and the evolution of thermal reaction norms. Trends Ecol. Evol. 18, 234–240 (2003).

34. J. Kingsolver, et al., Curve-thinking: Understanding reaction norms and developmental trajectories as traits 10.1002/9781118398814.ch3.

35. R. Izem, J. G. Kingsolver, Variation in continuous reaction norms: quantifying directions of biological interest. Am. Nat. 166, 277–289 (2005).

36. Y. Zhang, et al., Between- and within-population variations in thermal reaction norms of the coccolithophore Emiliania huxleyi. Limnol. Oceanogr. 59, 1570–1580 (2014).

37. G. M. Grimaud, F. Mairet, A. Sciandra, O. Bernard, Modeling the temperature effect on the specific growth rate of phytoplankton: a review. *Rev*. Environ. Sci. Technol. 16, 625–645 (2017).

38. C. T. Kremer, M. K. Thomas, E. Litchman, Temperature- and size-scaling of phytoplankton population growth rates: Reconciling the Eppley curve and the metabolic theory of ecology. Limnol. Oceanogr. 62, 1658–1670 (2017).

39. J. Norberg, Biodiversity and ecosystem functioning: A complex adaptive systems approach. Limnol. Oceanogr. 49, 1269–1277 (2004).

40. D. H. Janzen, Why Mountain Passes are Higher in the Tropics. Am. Nat. 101, 233–249 (1967).

41. K. S. Sheldon, R. B. Huey, M. Kaspari, N. J. Sanders, Fifty Years of Mountain Passes: A Perspective on Dan Janzen’s Classic Article. Am. Nat. 191, 553–565 (2018).

42. B. Chen, Patterns of thermal limits of phytoplankton. J. Plankton Res. 37, 285–292 (2015).

43. G. L. Wheeler, D. Sturm, G. Langer, *Gephyrocapsa huxleyi* (*Emiliania huxleyi*) as a model system for coccolithophore biology. J. Phycol. (2023) 10.1111/jpy.13404.

44. B. A. Read, et al., Pan genome of the phytoplankton Emiliania underpins its global distribution. Nature 499, 209–213 (2013).

45. S. Blanco-Ameijeiras, et al., Phenotypic Variability in the Coccolithophore Emiliania huxleyi. PLoS One 11, e0157697 (2016).

46. A. Rosas-Navarro, G. Langer, P. Ziveri, Temperature affects the morphology and calcification of Emiliania huxleyi strains. Biogeosciences 13, 2913–2926 (2016).

47. M. H. Conte, A. Thompson, D. Lesley, R. P. Harris, Genetic and Physiological Influences on the Alkenone/Alkenoate Versus Growth Temperature Relationship in Emiliania huxleyi and Gephyrocapsa Oceanica. Geochim. Cosmochim. Acta 62, 51–68 (1998).

48. S. R. Fielding, Emiliania huxleyi specific growth rate dependence on temperature. Limnol. Oceanogr. 58, 663–666 (2013).

49. P. von Dassow, P. V. Muñoz Farías, S. Pinon, E. Velasco-Senovilla, S. Anguita-Salinas, Do Differences in Latitudinal Distributions of Species and Organelle Haplotypes Reflect Thermal Reaction Norms Within the Emiliania/Gephyrocapsa Complex? Frontiers in Marine Science 8 (2021).

50. S. I. Anderson, T. A. Rynearson, Variability approaching the thermal limits can drive diatom community dynamics. Limnol. Oceanogr. 65, 1961–1973 (2020).

51. B. J. Kendrick, et al., Temperature-induced viral resistance in Emiliania huxleyi (Prymnesiophyceae). PLoS One 9, e112134 (2014).

52. R. M. Sheward, J. D. Liefer, A. J. Irwin, Z. V. Finkel, Elemental stoichiometry of the key calcifying marine phytoplankton Emiliania huxleyi under ocean climate change: A meta-analysis. Glob. Chang. Biol. 29, 4259–4278 (2023).

53. W. G. Sunda, S. A. Huntsman, Cobalt and zinc interreplacement in marine phytoplankton: Biological and geochemical implications. Limnol. Oceanogr. 40, 1404–1417 (1995).

54. R. F. Strzepek, P. W. Boyd, W. G. Sunda, Photosynthetic adaptation to low iron, light, and temperature in Southern Ocean phytoplankton. Proc. Natl. Acad. Sci. U. S. A. 116, 4388–4393 (2019).

55. P. Echeveste, P. Croot, P. von Dassow, Differences in the sensitivity to Cu and ligand production of coastal vs offshore strains of Emiliania huxleyi. Sci. Total Environ. 625, 1673–1680 (2018).

56. M. J. Frada, S. Keuter, G. Koplovitz, Y. Avrahami, Divergent fate of coccolithophores in a warming tropical ecosystem. Glob. Chang. Biol. 28, 1560–1568 (2022).

57. S. I. Anderson, A. D. Barton, S. Clayton, S. Dutkiewicz, T. A. Rynearson, Marine phytoplankton functional types exhibit diverse responses to thermal change. Nat. Commun. 12, 6413 (2021).

58. S. Rivero-Calle, A. Gnanadesikan, C. E. Del Castillo, W. M. Balch, S. D. Guikema, Multidecadal increase in North Atlantic coccolithophores and the potential role of rising CO₂. Science 350, 1533–1537 (2015).

59. K. M. Krumhardt, N. S. Lovenduski, N. M. Freeman, N. R. Bates, Apparent increase in coccolithophore abundance in the subtropical North Atlantic from 1990 to 2014. Biogeosciences 13, 1163–1177 (2016).

60. S. Dutkiewicz, et al., Dimensions of marine phytoplankton diversity. Biogeosciences 17, 609–634 (2020).

61. D. Demory, et al., Picoeukaryotes of the Micromonas genus: sentinels of a warming ocean. ISME J. 13, 132–146 (2019).

62. M. Ye, et al., Multi-trait analysis reveals large interspecific differences for phytoplankton in response to thermal change. Mar. Environ. Res. 188, 106008 (2023).

63. I. W. Bishop, S. I. Anderson, S. Collins, T. A. Rynearson, Thermal trait variation may buffer Southern Ocean phytoplankton from anthropogenic warming. Glob. Chang. Biol. 28, 5755–5767 (2022).

64. G. W. Gilchrist, Specialists and Generalists in Changing Environments. I. Fitness Landscapes of Thermal Sensitivity. Am. Nat. 146, 252–270 (1995).

65. M. J. Follows, S. Dutkiewicz, S. Grant, S. W. Chisholm, Emergent biogeography of microbial communities in a model ocean. Science 315, 1843–1846 (2007).

66. D. Tilman, Resources: A Graphical-Mechanistic Approach to Competition and Predation. Am. Nat. 116, 362–393 (1980).

67. X. Fu, J. Sun, Temperature driving vertical stratification regulates phytoplankton community structure in the Bohai Sea and Yellow Sea. Mar. Environ. Res. 194, 106320 (2024).

68. U. Sommer, “The Role of Competition for Resources in Phytoplankton Succession” in Plankton Ecology: Succession in Plankton Communities, U. Sommer, Ed. (Springer Berlin Heidelberg, 1989), pp. 57–106.

69. D. Tilman, S. S. Kilham, P. Kilham, Phytoplankton community ecology: The role of limiting nutrients. Annu. Rev. Ecol. Syst. 13, 349–372 (1982).

70. A. Toseland, et al., The impact of temperature on marine phytoplankton resource allocation and metabolism. Nat. Clim. Chang. 3, 979–984 (2013).

71. B. Andersson, et al., Intraspecific variation in metal tolerance modulate competition between two marine diatoms. ISME J. 16, 511–520 (2022).

72. W. G. Sunda, Trace metal interactions with marine phytoplankton. Biological oceanography (1989).

73. P. W. Boyd, et al., Marine Phytoplankton Temperature versus Growth Responses from Polar to Tropical Waters – Outcome of a Scientific Community-Wide Study. PLoS One 8, e63091 (2013).

74. D. R. Easterling, et al., Climate extremes: observations, modeling, and impacts. Science 289, 2068–2074 (2000).

75. G. A. Meehl, C. Tebaldi, More intense, more frequent, and longer lasting heat waves in the 21st century. Science 305, 994–997 (2004).

76. K. Mason-Jones, S. L. Robinson, G. F. C. Veen, S. Manzoni, W. H. van der Putten, Microbial storage and its implications for soil ecology. ISME J. 16, 617–629 (2022).

77. A. A. Malik, J. Puissant, T. Goodall, S. D. Allison, R. I. Griffiths, Soil microbial communities with greater investment in resource acquisition have lower growth yield. Soil Biol. Biochem. 132, 36–39 (2019).

78. S. Keuter, G. Koplovitz, A. Torfstein, M. J. Frada, Two-year seasonality (2017, 2018), export and long-term changes in coccolithophore communities in the subtropical ecosystem of the Gulf of Aqaba, Red Sea. Deep Sea Res. Part I 191, 103919 (2023).

79. W. F. Blankley, R. A. Lewin, Temperature responses of a coccolithophorid, Cricosphaera carterae, measured in a simple and inexpensive thermal-gradient device1. Limnol. Oceanogr. 21, 457–462 (1976).

80. C. J. Watras, S. W. Chisholm, D. M. Anderson, Regulation of growth in an estuarine clone of Gonyaulax tam arensis Lebour: Salinity-dependent temperature responses. J. Exp. Mar. Bio. Ecol. 62, 25–37 (1982).

81. J. P. Strock, S. Menden-Deuer, Temperature acclimation alters phytoplankton growth and production rates. Limnol. Oceanogr. 66, 740–752 (2021).

82. M. K. Thomas, C. T. Kremer, C. A. Klausmeier, E. Litchman, A global pattern of thermal adaptation in marine phytoplankton. Science 338, 1085–1088 (2012).

83. B. Bolker, M. B. Bolker, Package “bbmle.” Tools for General Maximum Likelihood Estimation 641 (2017).

84. S. Dutkiewicz, et al., Capturing optically important constituents and properties in a marine biogeochemical and ecosystem model. Biogeosci. Discuss. 12, 2607–2695 (2015).

85. B. Rudis, B. Bolker, J. Schulz, A. Kothari, J. Sidi, ggalt: extra coordinate systems, geoms’, statistical transformations, scales and fonts for “ggplot2” (2017).

86. H. Wickham, ggplot2: elegant graphics for data analysis.

87. R Core Team, R: A language and environment for statistical computing (2022) (September 22, 2023).

88. E. Neuwirth, R. C. Brewer, ColorBrewer palettes. R package version (2014).

89. K. C. Mundim, S. Baraldi, H. G. Machado, F. M. C. Vieira, Temperature coefficient (Q10) and its applications in biological systems: Beyond the Arrhenius theory. Ecol. Modell. 431, 109127 (2020).

90. A. E. F. Prowe, M. Pahlow, S. Dutkiewicz, A. Oschlies, How important is diversity for capturing environmental-change responses in ecosystem models? Biogeosciences 11, 3397–3407 (2014).

91. M. J. Brush, J. W. Brawley, S. W. Nixon, J. N. Kremer, Modeling phytoplankton production: problems with the Eppley curve and an empirical alternative. Mar. Ecol. Prog. Ser. 238, 31–45 (2002).

92. C. A. Stock, et al., Ocean biogeochemistry in GFDL’s earth system model 4.1 and its response to increasing atmospheric CO _2_. J. Adv. Model. Earth Syst. 12 (2020).

93. S. I. Anderson, et al., Phytoplankton thermal trait parameterization alters community structure and biogeochemical processes in a modeled ocean. Glob. Chang. Biol. 30 (2024).

94. K. F. Edwards, M. K. Thomas, C. A. Klausmeier, E. Litchman, Phytoplankton growth and the interaction of light and temperature: A synthesis at the species and community level. Limnol. Oceanogr. 61, 1232–1244 (2016).

95. S. Barton, G. Yvon-Durocher, Quantifying the temperature dependence of growth rate in marine phytoplankton within and across species. Limnol. Oceanogr. 64, 2081–2091 (2019).

96. E. Litchman, M. K. Thomas, Are we underestimating the ecological and evolutionary effects of warming? Interactions with other environmental drivers may increase species vulnerability to high temperatures. Oikos (2022) 10.1111/oik.09155.

97. K. M. Krumhardt, N. S. Lovenduski, M. D. Iglesias-Rodriguez, J. A. Kleypas, Coccolithophore growth and calcification in a changing ocean. Prog. Oceanogr. 159, 276–295 (2017).

98. E. Litchman, C. A. Klausmeier, Trait-Based Community Ecology of Phytoplankton. Annu. Rev. Ecol. Evol. Syst. 39, 615–639 (2008).

99. D. Atkinson, B. J. Ciotti, D. J. S. Montagnes, Protists decrease in size linearly with temperature: ca. 2.5% °C−1. Proceedings of the Royal Society of London. Series B: Biological Sciences 270, 2605–2611 (2003).

100. B. Chen, H. Liu, Relationships between phytoplankton growth and cell size in surface oceans: Interactive effects of temperature, nutrients, and grazing. Limnol. Oceanogr. 55, 965–972 (2010).

101. E. Marañón, Cell size as a key determinant of phytoplankton metabolism and community structure. Ann. Rev. Mar. Sci. 7, 241–264 (2015).

102. E. Marañón, et al., Unimodal size scaling of phytoplankton growth and the size dependence of nutrient uptake and use. Ecol. Lett. 16, 371–379 (2013).

103. S. G. Leles, N. M. Levine, Mechanistic constraints on the trade-off between photosynthesis and respiration in response to warming. Sci Adv 9, eadh8043 (2023).

104. Y. Fu, C. O’Kelly, M. Sieracki, D. L. Distel, Protistan grazing analysis by flow cytometry using prey labeled by in vivo expression of fluorescent proteins. Appl. Environ. Microbiol. 69, 6848–6855 (2003).

105. B. A. Ward, S. Dutkiewicz, O. Jahn, M. J. Follows, A size-structured food-web model for the global ocean. Limnol. Oceanogr. 57, 1877–1891 (2012).

106. F. M. Monteiro, et al., Why marine phytoplankton calcify. Sci Adv 2, e1501822 (2016).

107. C. T. Johns, et al., The mutual interplay between calcification and coccolithovirus infection. Environ. Microbiol. 21, 1896–1915 (2019).

