## Supplemental Text and Figures for "Intraspecific diversity in thermal performance determines phytoplankton ecological niche"

**Supplementary Information**

The Eppley curve and temperature variation background

Biological temperature response is generally predicted to be exponential (32) or to follow a power law relationship (89). Phytoplankton temperature response is often parameterized using the Eppley curve (27, 57, 82, 90), in which at increasing optimum temperatures, phytoplankton have exponentially increasing maximum growth rates. While the incompleteness of the Eppley curve has been widely documented (57, 91), it remains a concise way to represent the response of phytoplankton growth rates to temperature in ecosystem models. When a functional group is modeled as a single entity, individual thermal reaction norms are not resolved (82, 92). In the few ecosystem models that resolve variations in individual thermal response, thermal reaction norms are typically identically parameterized along the growth rate-optimum temperature continuum described by the Eppley curve, with each thermal reaction norm associated with a specific optimum temperature and a predetermined growth rate maximum (60, 65).

Coccolithophore diversity background

The “coccolithophore” functional type has been predicted to decrease in biomass with future climate warming due to its more slowly increasing growth rate with temperature (57, 93). Isolates available in culture are a biased subset of the total within-species diversity, and idealized yet inconsistent choices in factors like culture media, light levels, and culture vessels make different experiments difficult to appropriately compare. Firm control on differences between laboratory setups is necessary because thermal response is dependent on other ambient environmental conditions, and resource limitation may either enhance or eliminate observed thermal response (94–96). Further isolation of coccolithophores and thermal norm characterization is necessary to fully understand coccolithophore thermal biology. Many coccolithophore species (*e.g.*, genus *Umbellosphaera* and *Florisphaera profunda*) are abundant in the ocean but entirely missing from culture studies, which may leave significant gaps in our understanding of broader coccolithophore thermal ecology and future response to changing environmental temperature (97).

POC/PON and PIC measurements

PIC was determined by coulometric titration using a $CO_2$ coulometer (model CM5014, UIC Inc., Joliet, IL). Each sample was placed in a glass vial secured to a CM5130 acidification module and treated with 2 ml of 1N $H_3PO_4$ to liberate the $CO_2$ into a closed system to be measured.

In short, the coulometer contains a cell filled with a monoethanolamine solution and pH indicator and is placed between a light source and photodetector. Upon release $CO_2$ is adsorbed by the monoethanolamine to form a titratable acid causing the indicator to fade. The photodetector monitors the change of the color as a percent transmittance from the baseline. The amount of energy used to then neutralize the reaction using Faraday's Law allows for a calculation of $CO_2$ concentration (1 faraday of electricity results in the alteration of 1 gram equivalent weight of a substance during electrolysis).

<u>Strain-specific differences in the relationship of mean cell size and calcification to temperature</u>

Cell size is a key phytoplankton trait, as it controls nutrient affinity, population growth rate, and other traits via allometric scaling (38, 98). Cell size is thought to decrease with increasing temperature (99). However, questions about the phytoplankton cell size-temperature relationship persist, including why phytoplankton cell size may increase at very high temperatures, whether phytoplankton growth rate on a community scale decreases with increasing size, and whether the relationship between cell size and temperature is subject to adaptation (100–103). The impact of temperature on community allometry is determined by whether mean population size correlates with temperature and maximum growth rate within species. In our experiments, the estimated cell size of individual strains via flow cytometry (104) was differentially responsive to temperature. Most strains had decreasing cell size with temperature (Supplementary Figure 1C) up to the thermal optimum, followed by increasing cell size above the thermal optimum. Hence, mean cell size tended to track the growth rate of the culture as cell division rates increased. Some strains had relatively small changes in cell size (*e.g.*, strains RCC3963 and CCMP2090), which corresponded to higher values of the plateau parameter (Supplementary Figure 2A). Heterogeneity between strains in the responsiveness of their estimated cell size to temperature further supports the conclusion that different strains may use different mechanisms to respond to temperature under constant resource availability, which may be further investigated using molecular techniques. In some functional group modeling efforts, larger cell size phytoplankton types have lower maximum growth rate at the same thermal optimum value (60, 105). We did not find a correlation between maximum growth rate and mean cell size. The coexistence of strains with lower growth rates at the same thermal optimum suggests either that diversity in maximum growth rates in a population is valuable, or that the mechanisms that result in lower maximum growth rates at a single temperature are not determined solely by allometry and cannot necessarily be predicted by mean cell size.

Calcification is an important trait among coccolithophores (97, 106). While the precise reason that marine phytoplankton calcify is still a subject of investigation (106), calcification has a confounding influence on other aspects of coccolithophore environmental response, for example that nutrient limitation appears to simultaneously increase calcification rate and decrease coccolithophore growth rate (97). Calcification varies significantly among coccolithophore taxa, particularly within the *Gephyrocapsa* species complex, strains of which have among both the highest and lowest relative rates of calcification (97). Strains of *G. huxleyi* in culture collections may or may not calcify, and as such, five of the twelve strains we used in our experiments are noted by their respective culture collections to be naked (strains RCC874, CCMP375, CCMP379, CCMP2090, and CCMP374; Roscoff Culture Collection and Bigelow National Center for Marine Algae and Microbiota). Cell complexity can be measured via flow cytometry using side scatter, which has previously been established as a useful tool for estimating calcification in *G. huxleyi (107)*. We found that alongside cell size, calcification extent via side scatter also tended to decrease with temperature among calcified strains (Supplementary Figures 1, 12, 13), which we corroborated for two strains using particulate inorganic carbon measurements (Supplementary Figure 15; Supplementary Table 3). This supports previous observations that coccolithophore calcification decreases with rising temperatures (97).

Model output re-analysis from Dutkiewicz et al. (2020) diversity model

In Dutkiewicz et al. (2020), a different model parameterization, which we refer to as the “Darwin parameterization” was used to modulate temperature impact on thermal response with identical parameters across the range of thermal reaction norms which were simulated:

$$\gamma_j^T = \tau_T exp\left(A_T\left(\frac{1}{T} - \frac{1}{T_n}\right)\right)exp\left(-B_T\left|T - T_{oj}\right|^b\right) \quad (2)$$

Wherein the parameter $\tau_T$ is used for scaling, $T_{oj}$ was varied according to temperature optimum, and the other parameters are shape parameters which were held constant across the 10 thermal types simulated in the model. The equation for the growth rate of the coccolithophore functional type is:

$$\mu_j = P_{mj}^C\left(1 - exp\left(\frac{-\Lambda_{E_j\theta_j}}{P_{mj}^C}\right)\right)$$

in the Darwin model, where $P_{mj}^C$ is the maximum light-saturated photosynthesis rate (1.03 for coccolithophores), $\Lambda_{Ej}$ is the slope of the photosynthesis-irradiance curve multiplied by irradiance (40 for coccolithophores), and $\theta_j$ is the chlorophyll *a* to carbon ratio (0.2 for coccolithophores) (84). Because the Darwin model includes an additional scaling factor for growth rate on the basis of cell size:

$$a * \left(\frac{4}{3}\pi r^3\right)^b$$

Using the values $a = 2.1$ and $b =- 0.15$ for coccolithophores, a constant scaling factor of 1.18 was added to the growth rate model above for a coccolithophore of ESD 4.5μm.

We used model output for coccolithophores that were derived from the temperature response curve with the following parameter values from (60) ( $\tau_T = 0.8$, $A_T =- 4000$, $T_N = 293.15$, $B_T = 3 \times 10^{-4}$, and $b = 4$) for the Arrhenius equation.

We evaluated whether a thermal diversity-resolving model reflected overlap in the thermal niche of coccolithophores, or whether the model parameterization implies strong geographic differentiation between thermal types. The relationship between local model environmental temperature and thermal type survivability was not explored in the original study. To do this, we reanalyzed a year of model output from the 2020 simulation collected at third-daily resolution (one measurement every 3 days) (60). We evaluated how daily coccolithophore biomass varies by thermal type, which had not been addressed in the previous study, and also calculated the average deviation of the local environmental temperature from the thermal optimum of the type. This enabled exploration of how tightly coupled thermal optimum is to local temperature in a simulation where many linearly-spaced thermal types are represented. If the maximum growth rate of the coccolithophores needs to be above some threshold in order for them to be successful in the ecosystem model, the varying shape of the thermal curve of the acclimated strains of coccolithophores that we reared in the lab study may sometimes underlie a competitive advantage in flexibility in response to temperature.

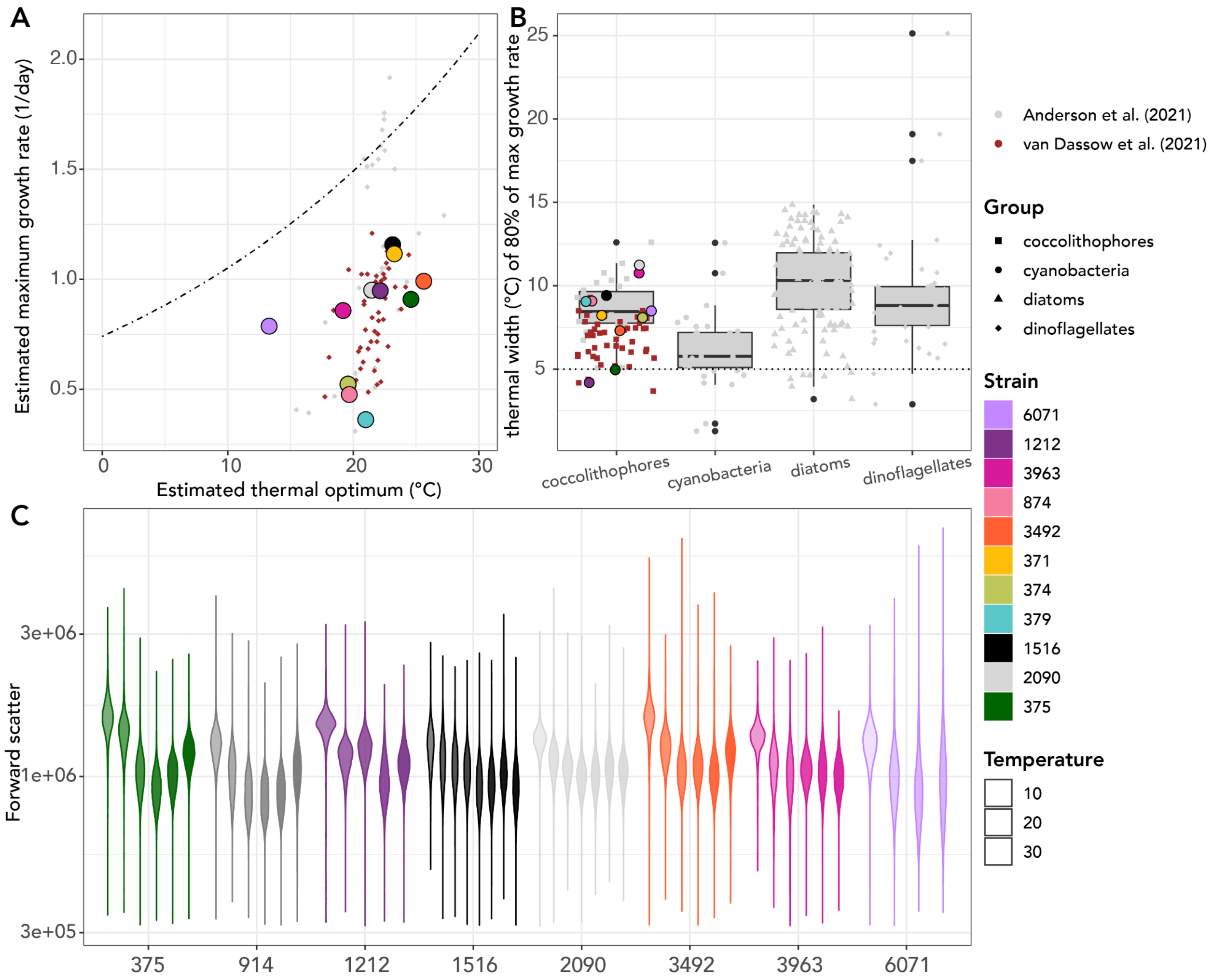


**Supplementary Figure 1. Maximum growth rates vary despite little overall variability in cell size.** A: Comparison of estimated maximum growth rates and thermal optima of the 12 strains of *Gephyrocapsa huxleyi* tested in the context of two other recent studies. The dotted line above the points identifies the Eppley relationship fit to coccolithophores from Anderson *et al*. (2021). B: "Plateau parameter" for tested strains of *Gephyrocapsa huxleyi* as compared to other coccolithophores and functional groups. This parameter is calculated as the width of temperatures at which growth rate is within 80% of the maximum growth rate recorded; strains with higher values for the plateau parameter have a wider range of near-optimum temperatures at which growth rates fall into a similar range as compared to the strain's maximum growth rate.C: Cell size as estimated from the forward scatter on the flow cytometer. Different temperatures tested are shown from left to right within each strain group. The smallest cell size was found close to each strain's thermal optimum, but overall cell sizes were not significantly different between strains.

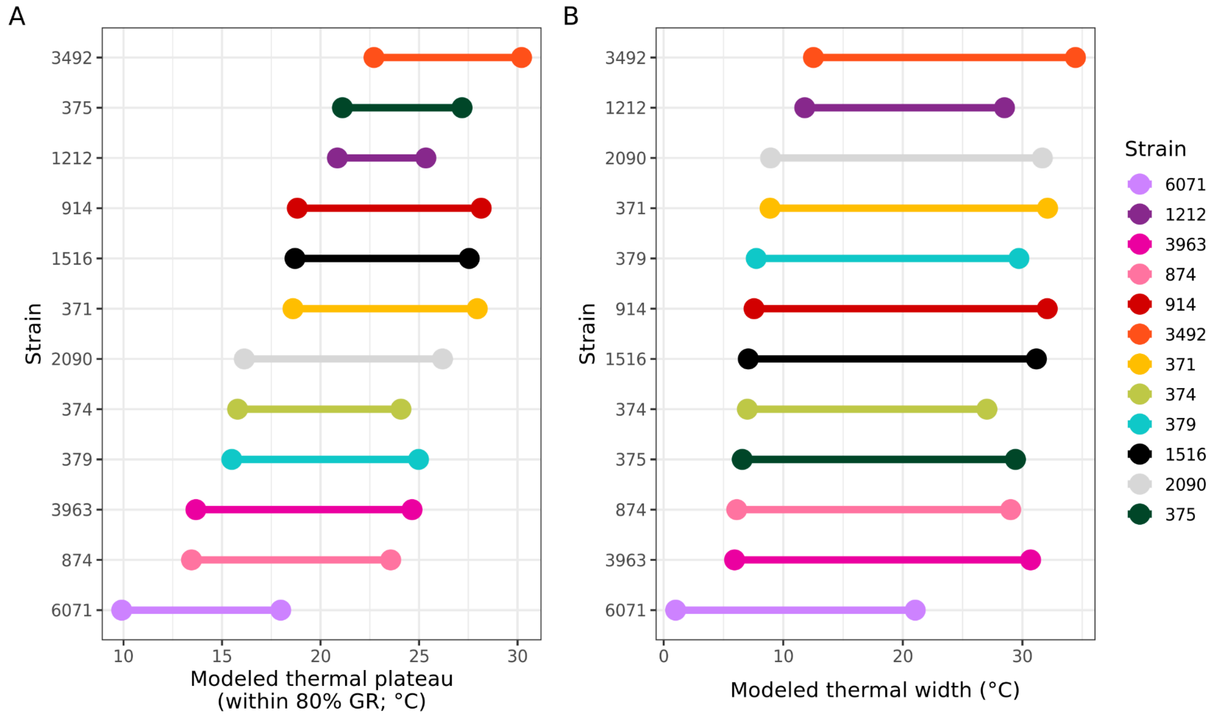


**Supplementary Figure 2. Comparison of thermal plateau (temperatures within 80% of growth rate) (A) vs. thermal width (temperatures with nonzero growth rate) in the modeled Norberg curves (B) for the 12 strains.** A wider thermal plateau indicates that more temperatures are survivable and also have relatively high growth rates. Some strains, e.g., RCC1212, had much narrower modeled thermal niche widths than others, e.g., CCMP2090.

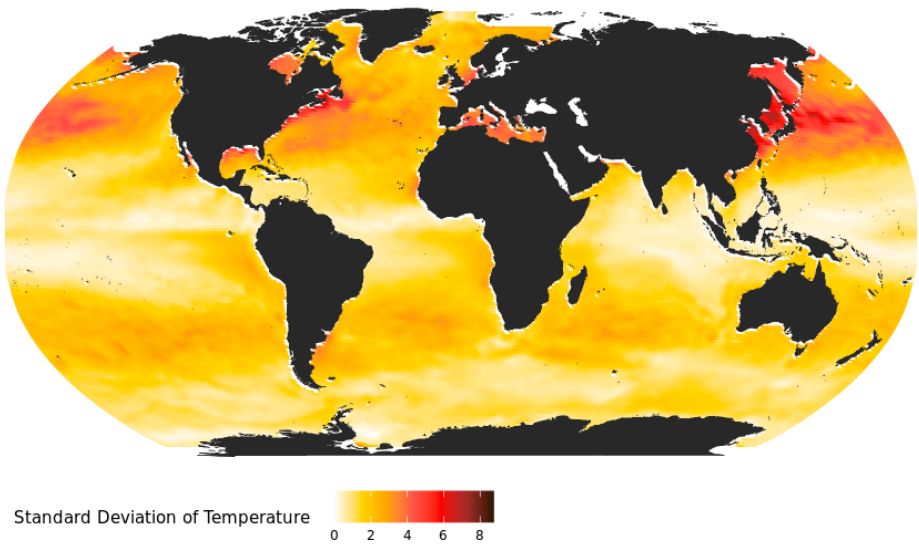


**Supplementary Figure 3.** Standard deviation of local water temperature at each grid point in Darwin model simulation using 1-month-averaged temperature resolution.

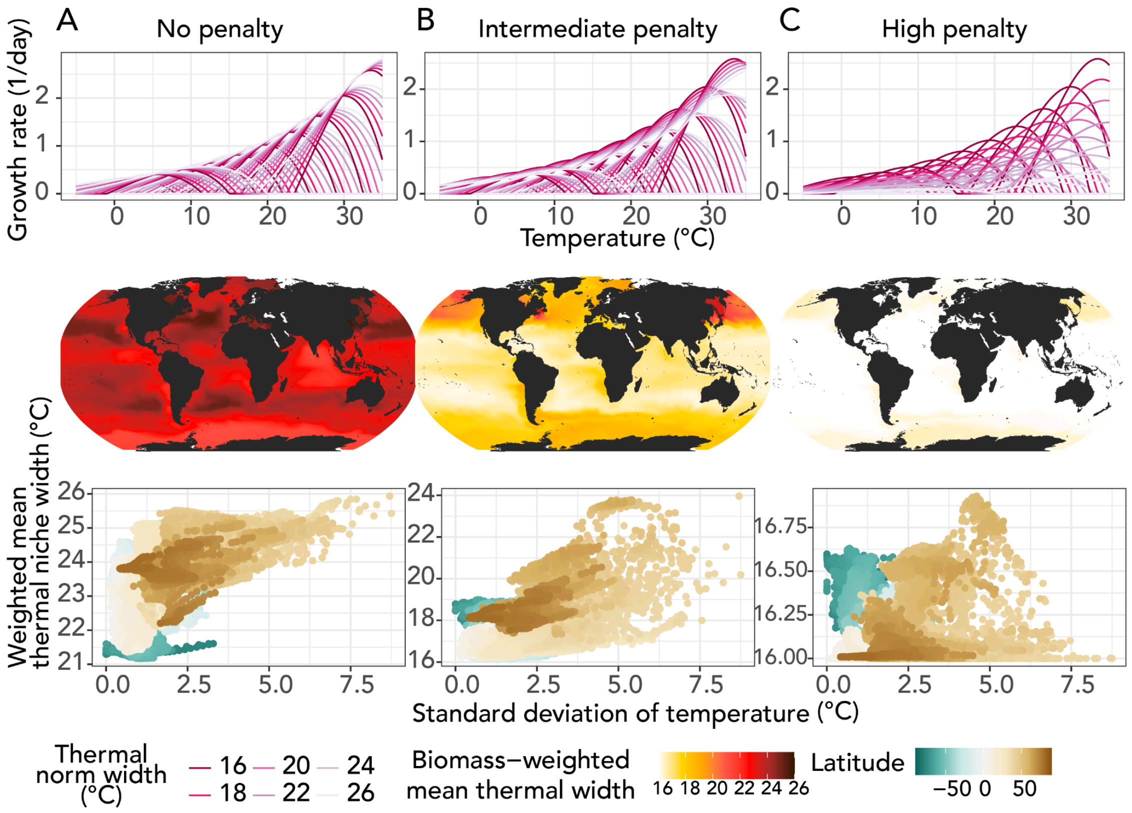


**Supplementary Figure 4**. Comparison of three tested penalties on growth rate with increasing thermal width. Each column of the figure shows (top) the growth curves of the different thermal optima and thermal niche widths of the 60 phytoplankton types included in each simulation, (middle) a map of biomass-weighted mean thermal width for the final year of the simulation, and (top) the standard deviation of monthly temperature at each latitude point (indicated by color) compared to the weighted thermal niche width. In (A) without any penalty, all parameters except thermal niche width are kept constant; generalists dominate, but weighted mean thermal niche width correlates with temperature variation. In (B) with the intermediate penalty retained for the final study, thermal niche width and thermal variability correlate to some extent and a greater diversity of niche widths reach high biomass. In (C), a very significant penalty results in specialists dominating, but with no correlation to local temperature variability.

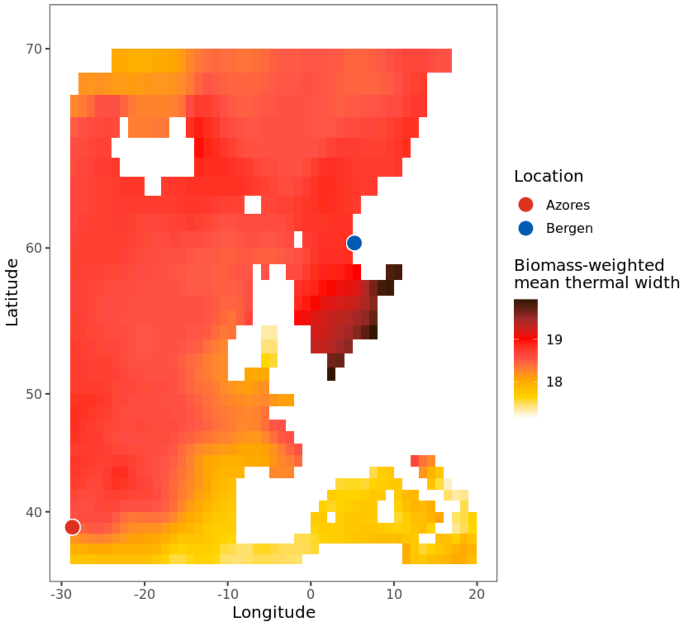


**Supplementary Figure 5.** Plot of mean thermal niche widths at each latitude-longitude location around Bergen, Norway and Azores, Portugal, demonstrating that Azores is located around a model front between generalist and specialist abundance.

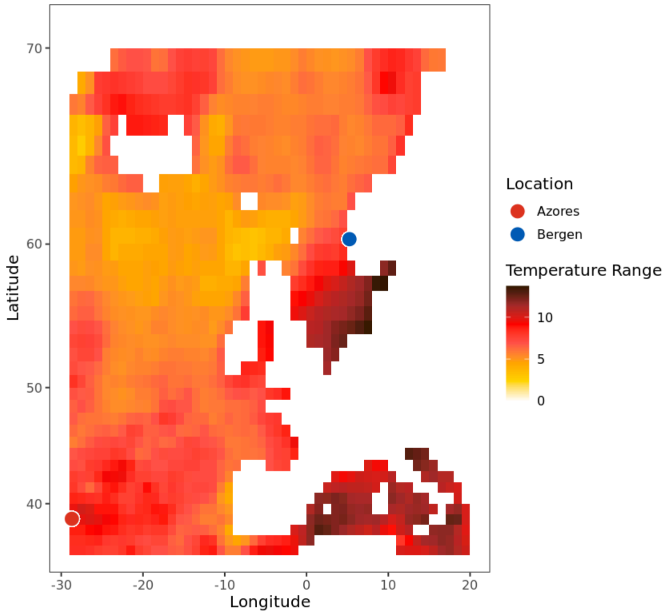


**Supplementary Figure 6.** Temperature range in the region of the Darwin model that included both Azores, Portugal and Bergen, Norway. The location of the isolation of laboratory *G. huxleyi* at each location is indicated by a filled circle on the map.

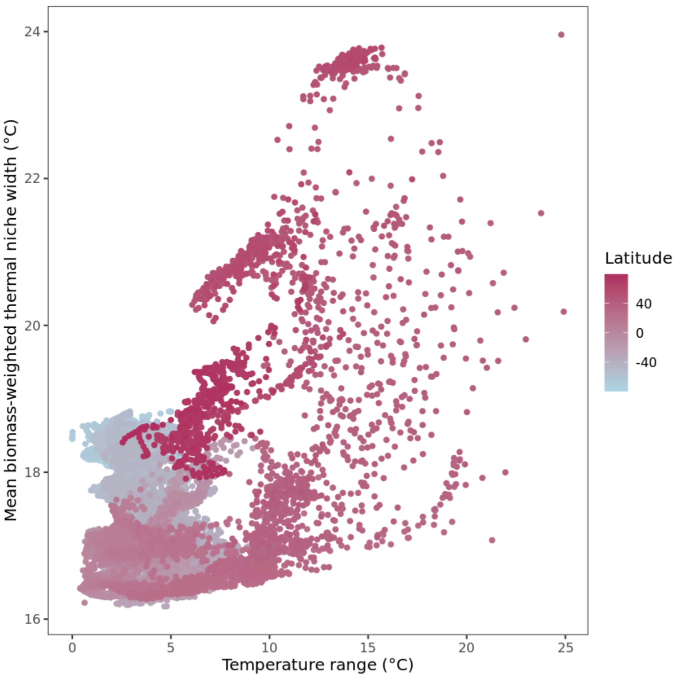


**Supplementary Figure 7.** Temperature range (range of temperatures experienced at grid point after 1-monthly averaging; x-axis) compared to the biomass-weighted thermal niche width at that location. Some, but not clear, correlation between thermal variability and biomass-weighted niche width.

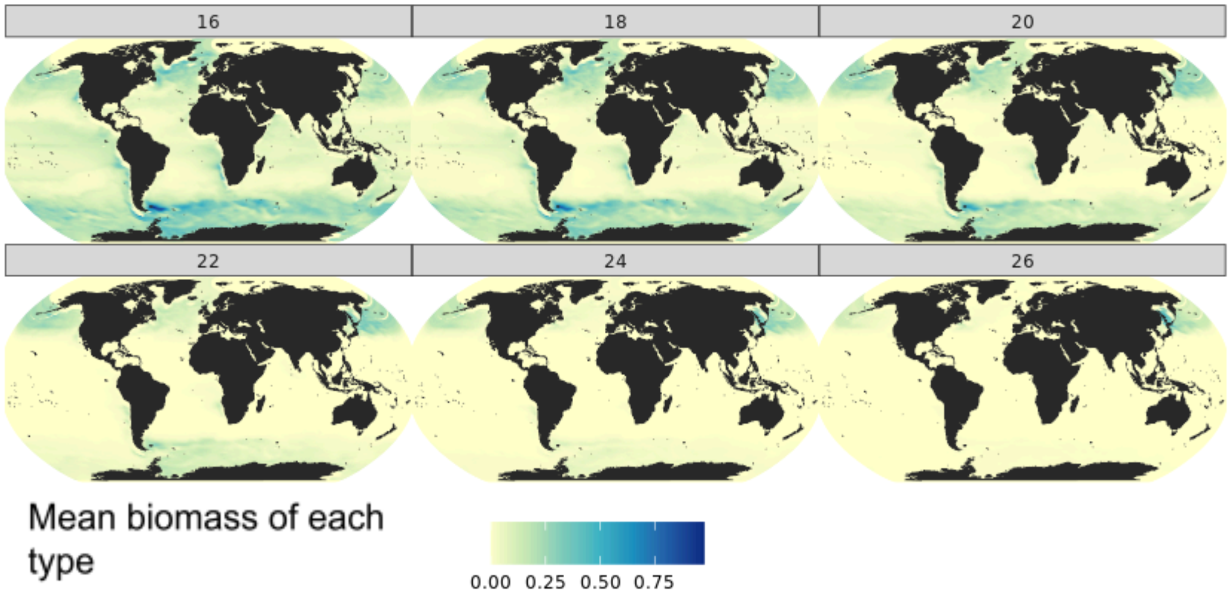


**Supplementary Figure 8.** Geographic extent of the 6 different thermal niche widths included in the generalist-specialist simulation. Fill indicates the amount of carbon biomass while facets indicate each of the 6 different thermal niche widths.

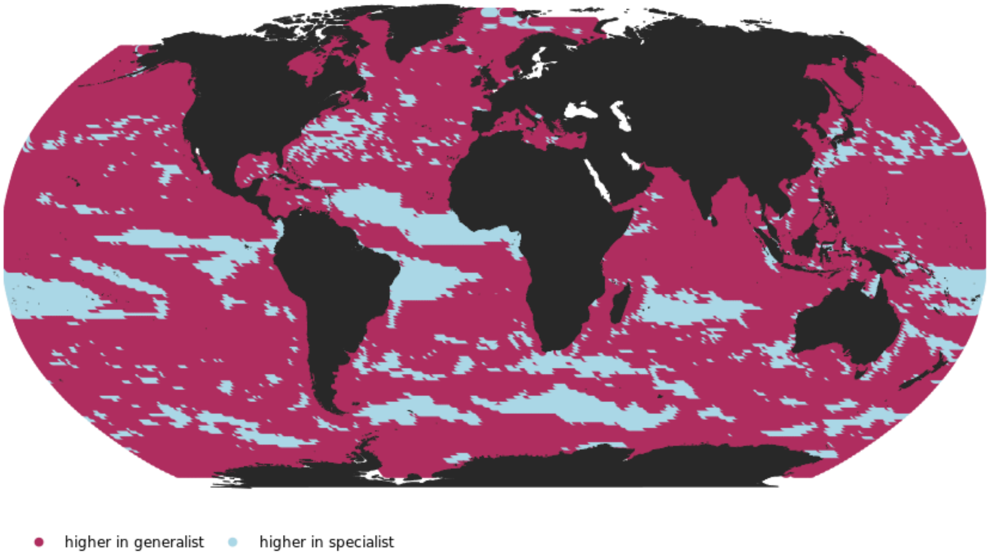


**Supplementary Figure 9.** Comparison of effect between the specialist-only or generalist-only simulation; grid points colored red had higher biomass in the generalist-only simulation, whereas grid points colored blue had higher biomass in the specialist-only simulation.

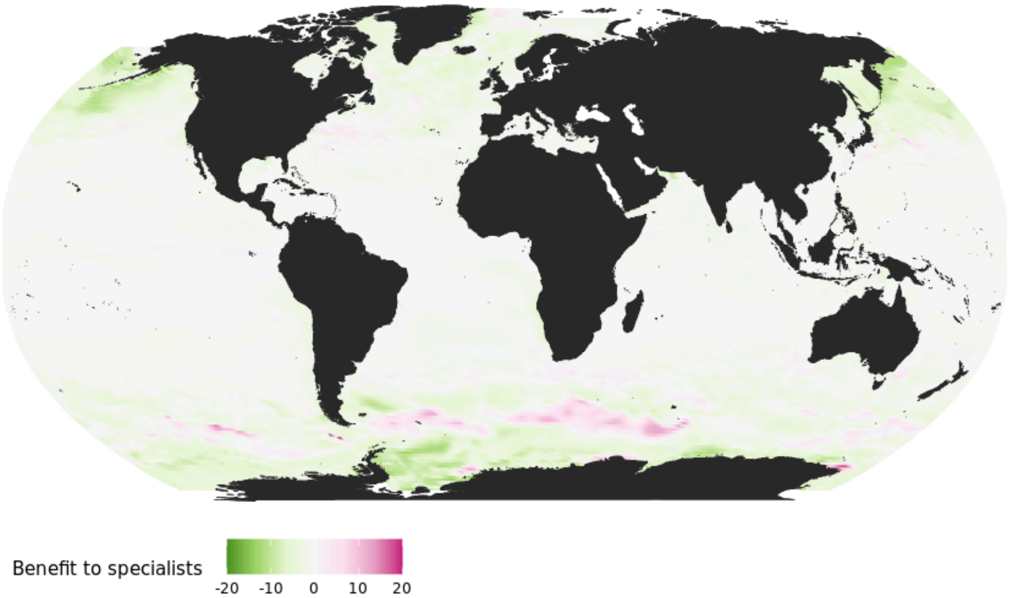


**Supplementary Figure 10.** Percent difference in biomass between the generalist-only and specialist-only simulations. Positive percentage difference numbers indicate higher biomass in the specialist-only simulation, whereas negative percentage difference numbers indicate higher biomass in the generalist-only simulation.

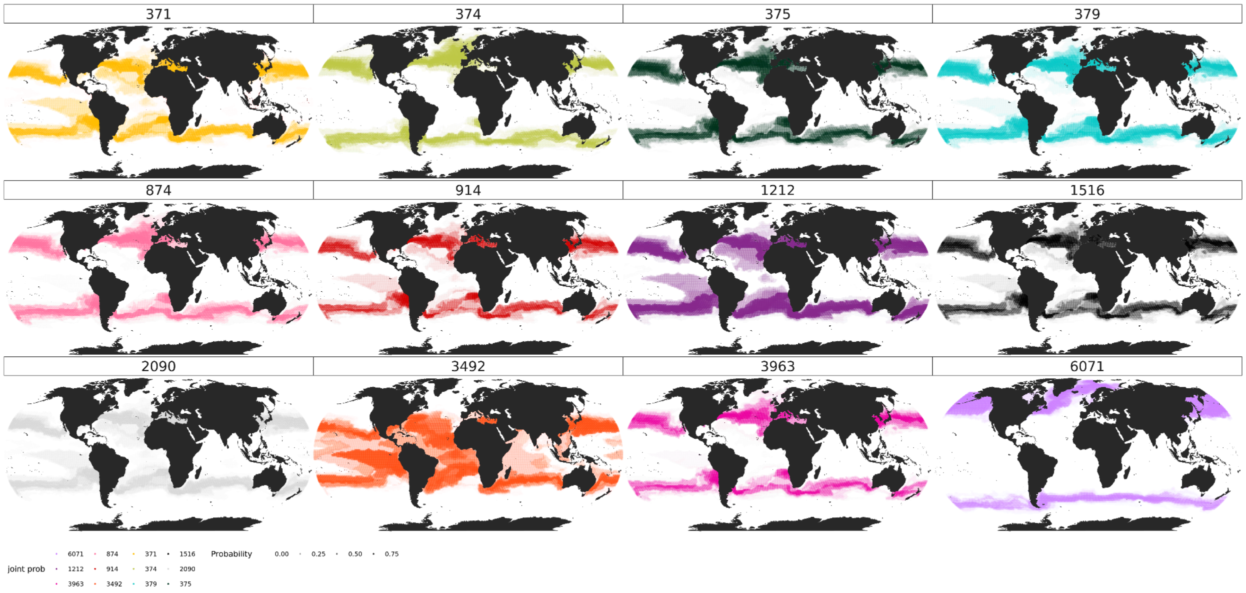


**Supplementary Figure 11.** Predicted geographic extent of all strains in the ecosystem model based on the probability calculation workflow.

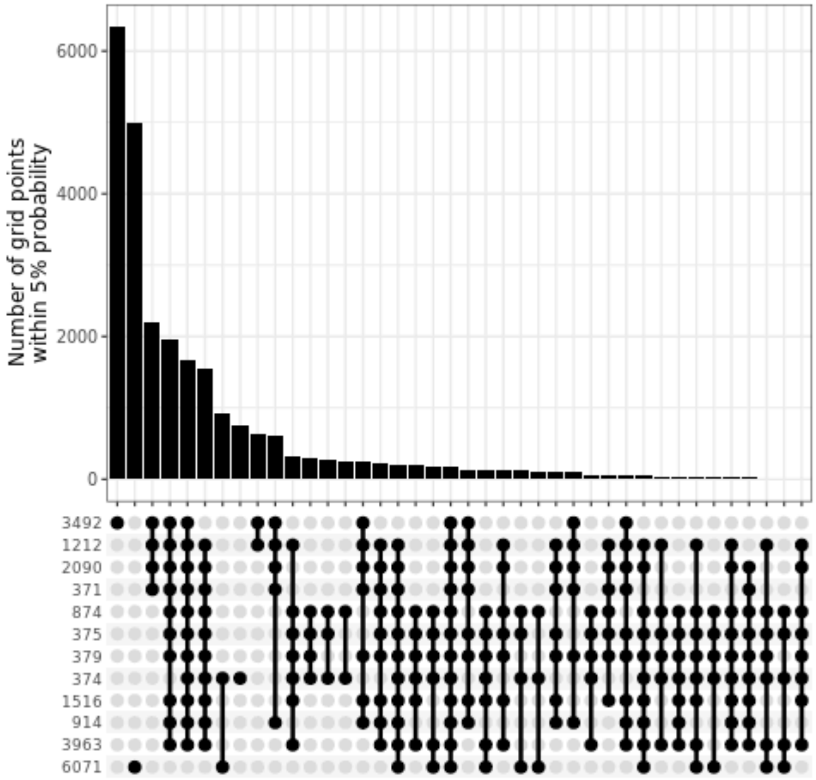


**Supplementary Figure 12.** Overlapping geographic locations (grid points) of the 12 strains, indicating that RCC3492 and RCC6071 on the far extremes of thermal range have the most gridded locations uniquely expected to be inhabited by those strains.

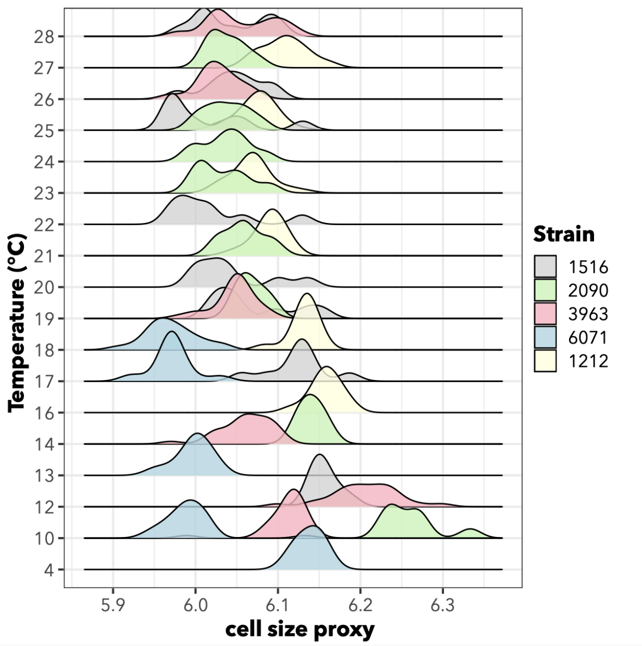


**Supplementary Figure 13.** Forward scatter (cell size proxy; x-axis) measured for each of 5 strains (fill of distributions) at a variety of temperatures (y-axis). Higher forward scatter approximately indicates larger cell size for measured cells.

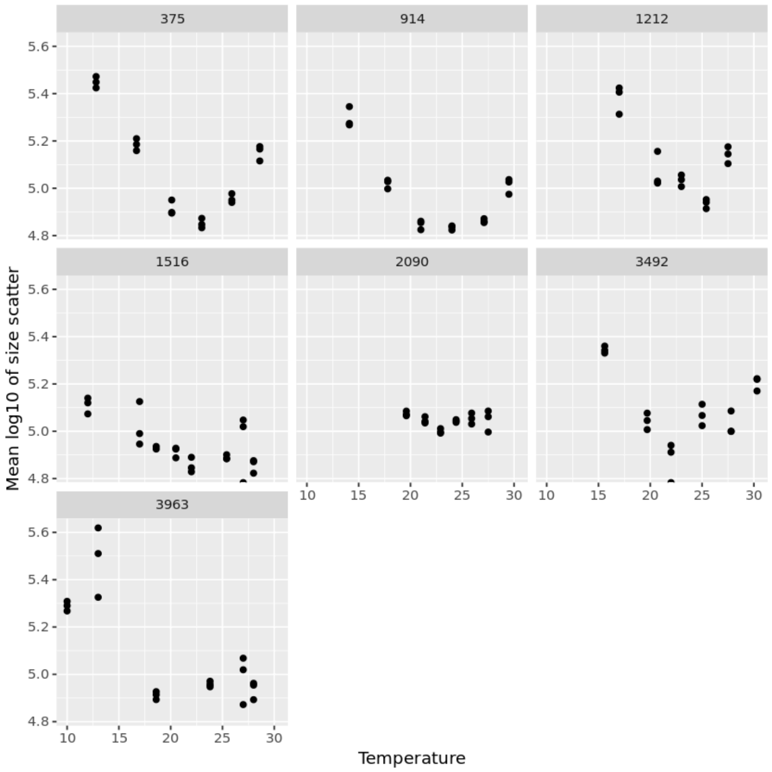

**Supplementary Figure 14.** Size scatter from flow cytometry measurements for seven strains of *G. huxleyi*. Side scatter can be used as a cell complexity or calcification proxy. In this figure, higher size scatter is expected to correspond to higher calcification in strains that calcify.

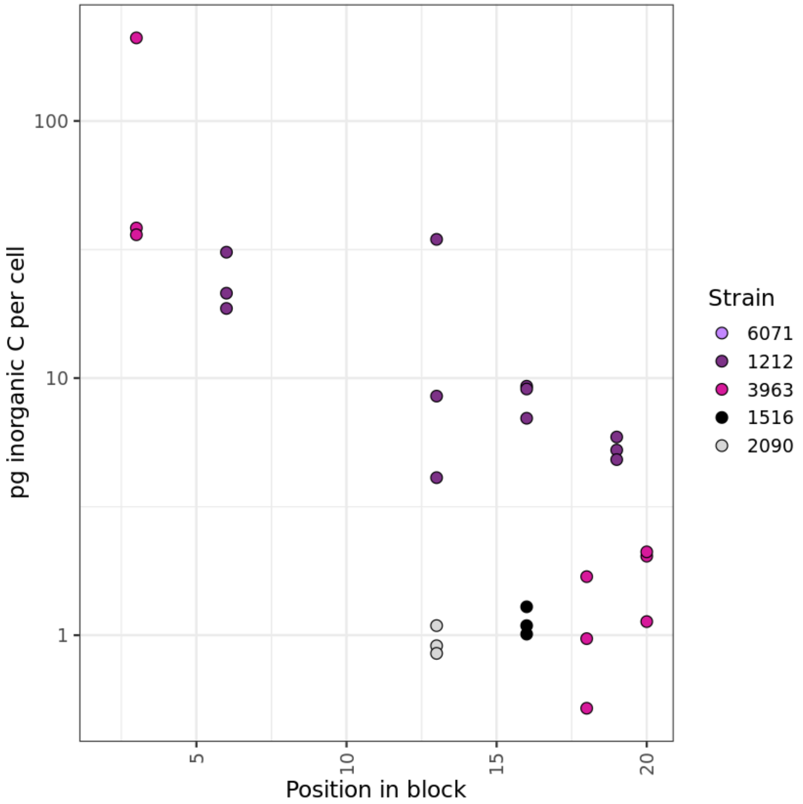


**Supplementary Figure 15.** Particulate inorganic carbon measurements for five of the twelve strains. Higher measured particulate inorganic carbon indicates a higher rate of calcification. Not all measured strains were calcifying, but those that did calcify appeared to calcify the most at lower temperatures.

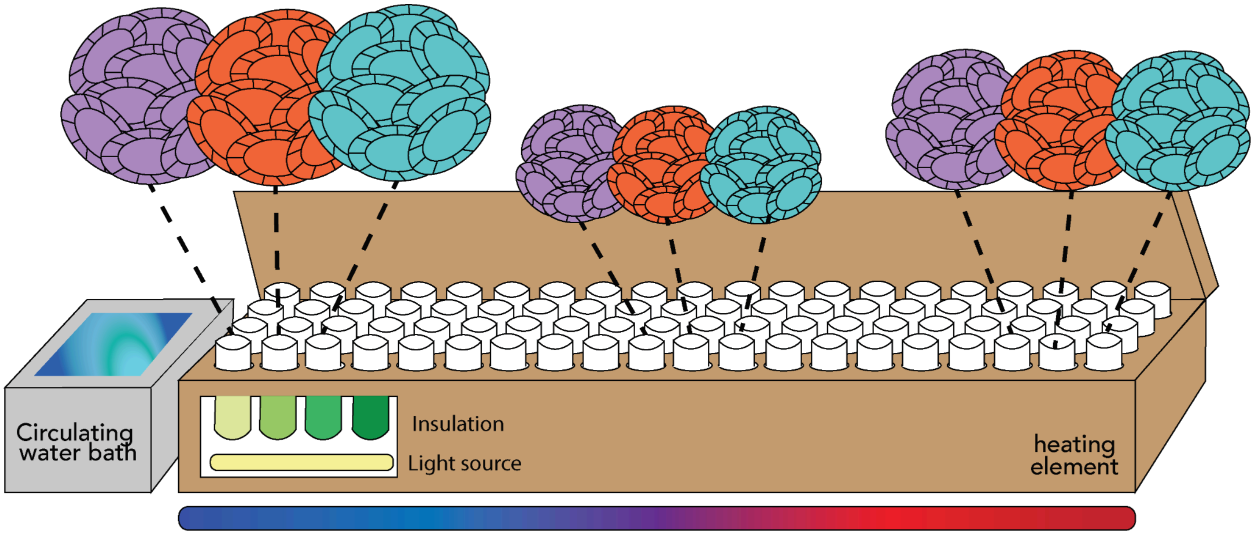

**Supplementary Figure 16.** Diagram of thermal block experimental setup for thermal acclimation experiments in the laboratory for twelve strains of *G. huxleyi*.

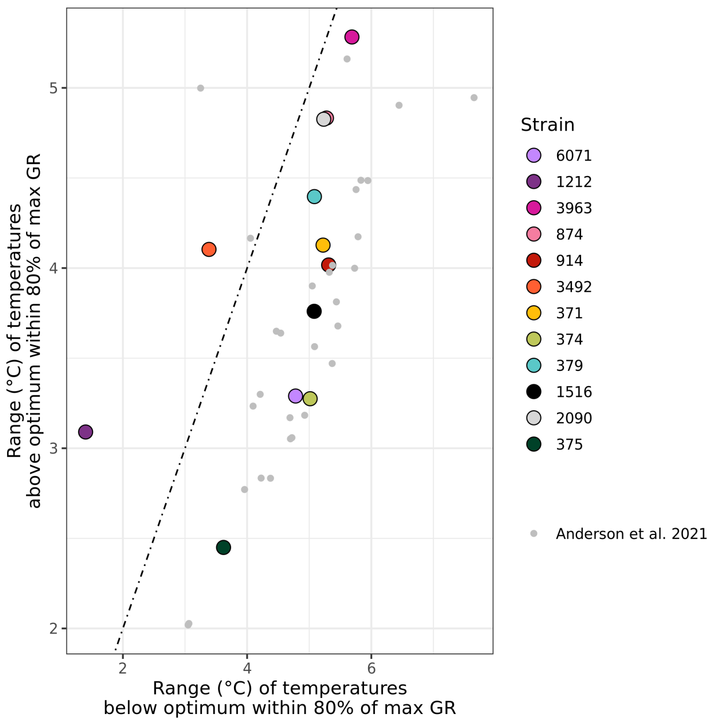


**Supplementary Figure 17.** Range of temperatures above vs. below maximum growth rate. The range of temperatures within 80% of the maximum measured growth rate and above the thermal optimum (y-axis) as compared to the range of temperatures within 80% of the measured maximum growth rate and below the thermal optimum (x-axis). The dotted line indicates a one-to-one relationship, hence points above this line have more survivable temperatures above the thermal optimum, and points below the line have more survivable temperatures below the thermal optimum.

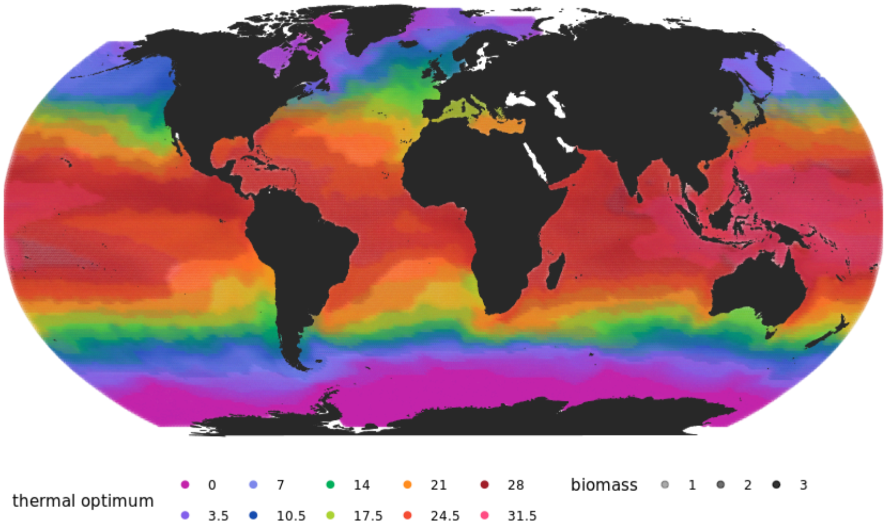

**Supplementary Figure 18.** Overlapping geographic ranges of the 10 thermal optimum values used in the diversity-resolving simulation. Higher alpha value (lower transparency) indicates higher biomass.

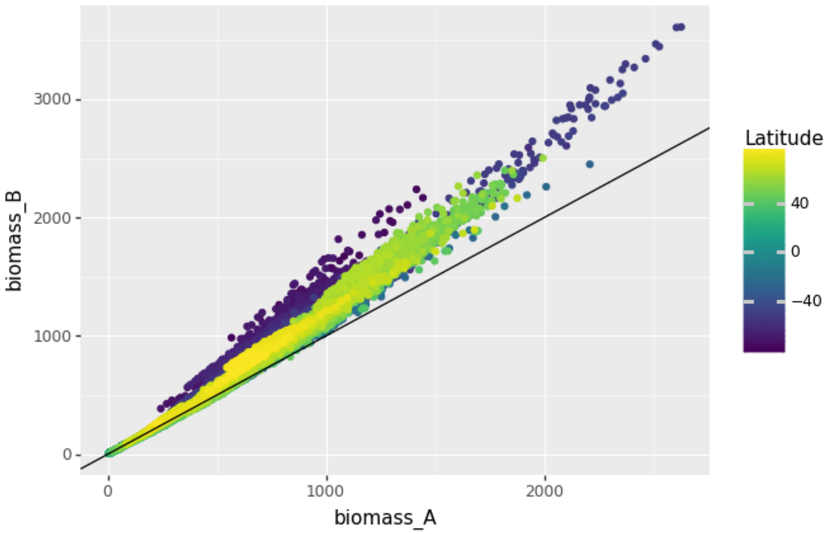


**Supplementary Figure 19.** Comparison of summed biomass in the specialist-only simulation (x-axis) to the summed biomass at each latitude-longitude point in the generalist-specialist simulation (y-axis). The solid black line indicates the one-to-one line.

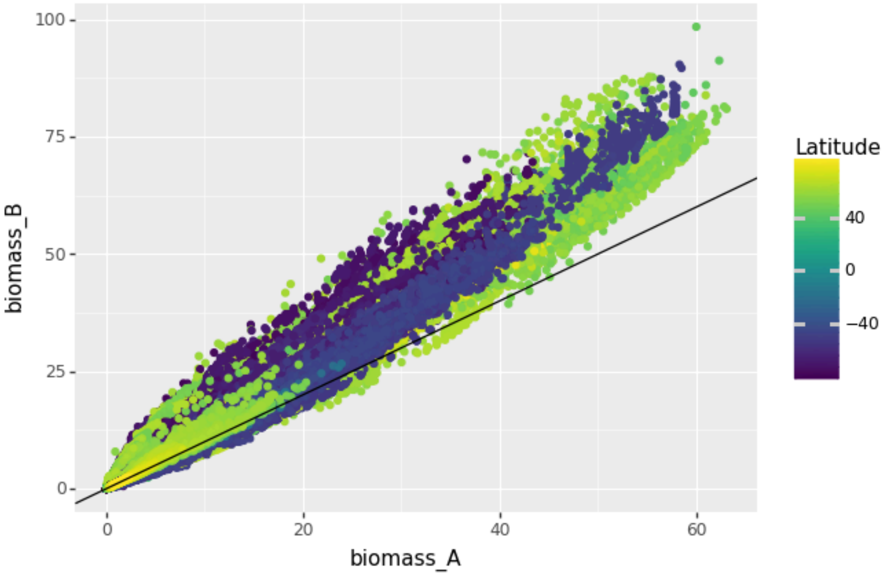


**Supplementary Figure 20.** Comparison of summed biomass in the specialist-only simulation (x-axis) to the summed biomass at each latitude-longitude-time (time in months) point in the generalist-specialist simulation (y-axis). The difference between this Supplementary Figure and the previous one shows that phenology was also shifted by the addition of generalists to the model, in addition to overall biomass.

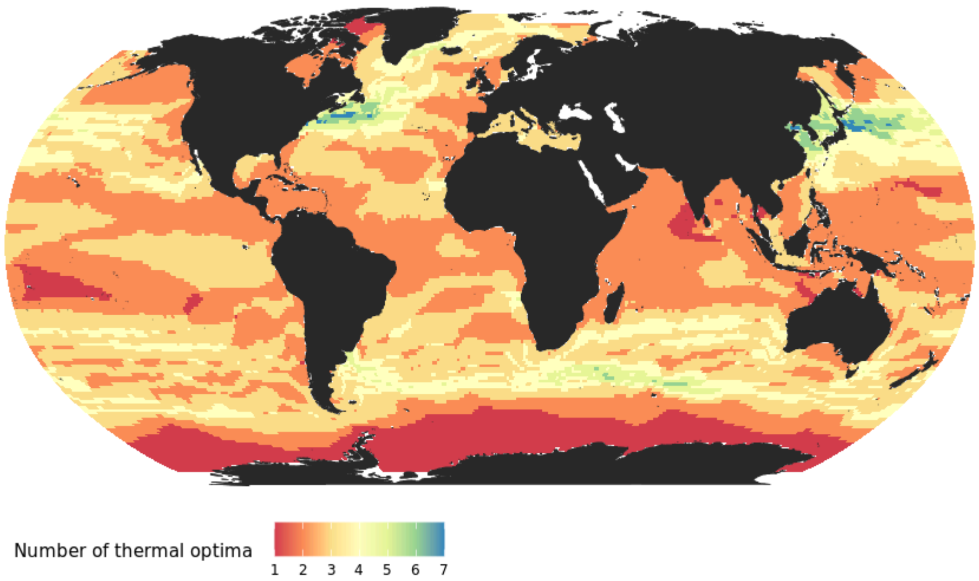


**Supplementary Figure 21.** Number of coexisting thermal optimum values at each model grid point when the generalist-specialist simulation was used.

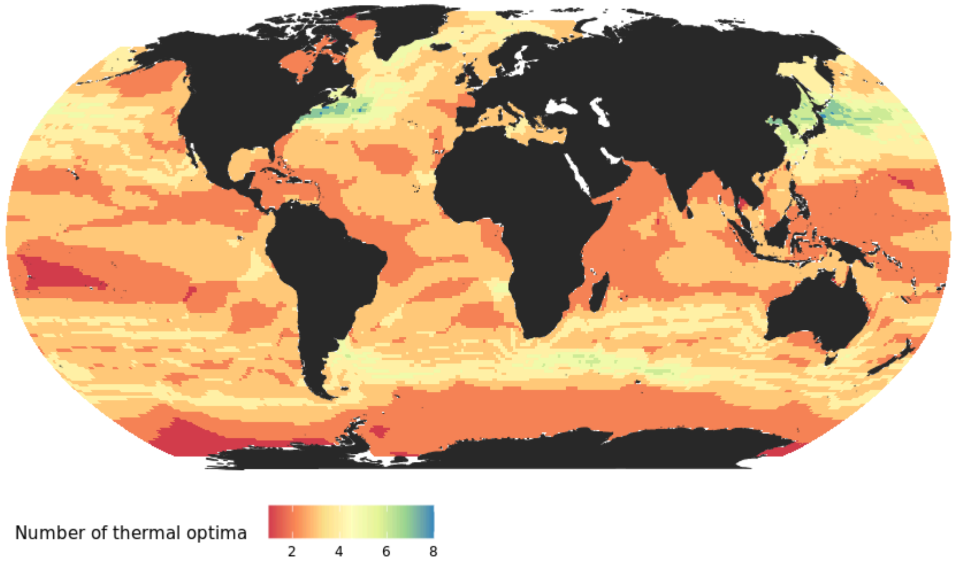


**Supplementary Figure 22.** Number of coexisting thermal optimum values at each model grid point when the specialist-only simulation was used.

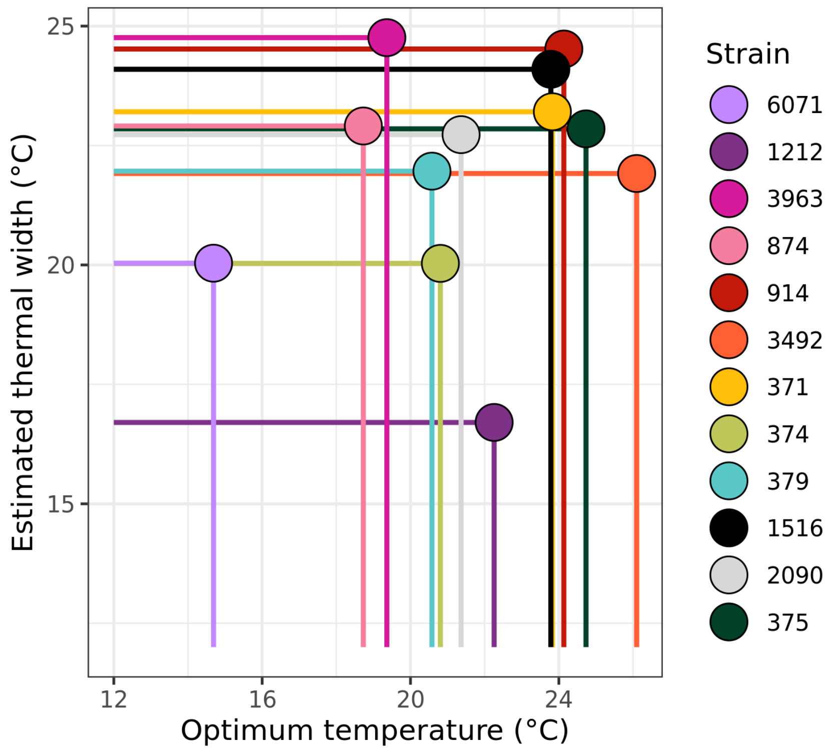


**Supplementary Figure 23.** Comparison of estimated thermal width (y-axis) and temperature optimum value (x-axis) for the 12 tested strains.

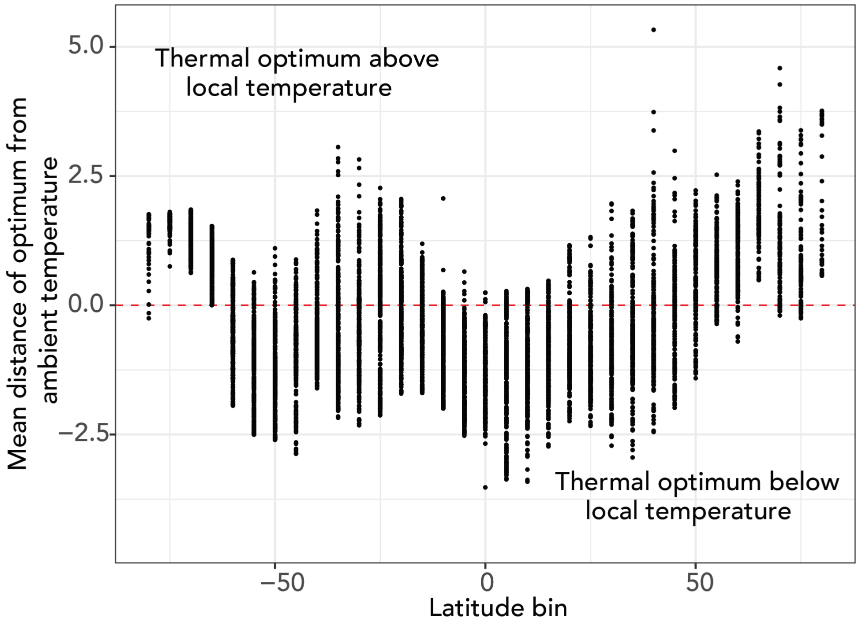


**Supplementary Figure 24.** Mean difference between optimum temperature and local monthly environmental temperature among strains present with at least 1% biomass vs. binned latitude coordinates. A positive distance indicates that thermal optimum value of present strains tended to be above the local environmental temperature, whereas a negative difference indicates that thermal optimum tended to be below water temperature.

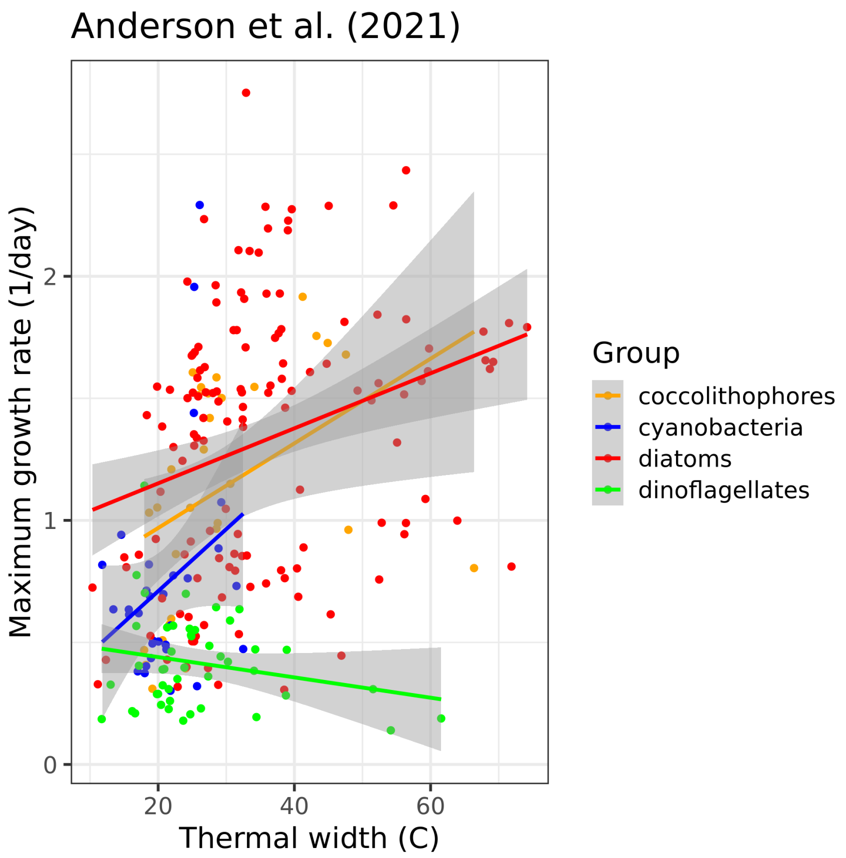


**Supplementary Figure 25.** Comparison of thermal niche width (degrees Celsius; x-axis) and maximum group rate (1/day; y-axis) for all of the compiled data from Anderson *et al*. (2021) (57). No functional type had a significant decline in maximum growth rate with increasing thermal width, as expected based on theory.

**Supplementary Table 1. Growth rate data from experiments - supplied as file**

**Supplementary Table 2. Thermal parameters calculated for each strain - supplied as file**

**Supplementary Table 3. Particulate inorganic carbon - supplied as file**
